# Resynthesis: Marker-based partial reconstruction of elite genotypes in clonally-reproducing plant species

**DOI:** 10.1101/2020.05.09.085738

**Authors:** Iban Eduardo, Simó Alegre, Konstantinos G. Alexiou, Pere Arús

## Abstract

We propose a method for marker-based selection of cultivars of clonally-reproducing plant species which keeps the basic genetic architecture of a top-performing cultivar (usually a partly heterozygous genotype), with some agronomically relevant differences (such as production time, product appearance or quality), providing added value to the product or cultivation process. The method is based on selecting a) two complementary nearly-inbred lines from successive selfing generations (ideally only F2 and F3) of large size, that may generate individuals with most of their genome identical to the original cultivar but being homozygous for either of the two component haplotypes in the rest, and b) individuals with such characteristics already occurring in the F_2_. Option a) allows for introgressing genes from other individuals in one or both of these nearly-inbred lines.

Peach, a woody-perennial, clonally-reproduced species, was chosen as a model for a proof of concept of the Resynthesis process due to its biological characteristics: self-compatibility, compact and genetically well-known genome, low recombination rates and relatively short intergeneration time (3-4 years). From 416 F_2_ seedlings from cultivar Sweet Dream (SD), we obtained seven individuals with 76-94% identity with SD, and selected five pairs of complementary lines with average homozygosity of the two parents ≥0.70 such that crossing would produce some individuals highly similar to SD. The application of this scheme to other species with more complex genomes or biological features, including its generalization to F1 hybrids, is discussed.

## INTRODUCTION

A few high-quality varieties dominate the market of many clonally propagated species. Reproducing the genotype of a top performing cultivar with a minor but significant improvement may have a high market value (e.g., a change in bloom or maturity time, skin or flesh fruit color, flesh sugar or acidity content and composition, and postharvest behavior). With classical approaches (sexual crossing and progeny selection) this is extremely unlikely, and to our knowledge, it has never been tried. Other approaches have difficult-to-overcome regulatory issues and sometimes a negative social perception (genetically modified or gene-edited materials), or are rare and occur unpredictably (somatic mutations). Resynthesis is an approach based on conventional breeding technologies that, with the help of molecular markers, allows the selection of two homozygous (or nearly homozygous) complementary individuals that, when crossed, produce offspring with a similar, albeit different, genetic composition from the original cultivar. Homozygous fragments of either of the two parents in specific genome regions that were heterozygous in the original cultivar could result in phenotypic changes for traits of interest in the context of the performance of a well-defined variety. In addition, the development of two parental lines allows for the incorporation of an exotic gene (disease resistance or fruit quality trait) in one of them and the Resynthesis of the improved cultivar in a predictable time frame (two generations after the first cross).

The number of crossovers per chromosome and meiosis is limited, usually between one and three in plants (Mercier et al. 2015), and the frequency of non-recombined chromosomes is high enough to find certain specific combinations (fully homozygous or heterozygous chromosomes) in a selfed offspring of reasonable size (Koumproglou et al. 2002). The basis of Resynthesis is that, with the help of markers covering the whole genome, it is possible to select fully homozygous and complementary genotypes to reproduce (resynthesize) an initial partly heterozygous genotype (i.e., a cultivar) with a limited number of selfing or backcross generations. The “Reverse Breeding” method (Dirks et al. 2009) is a similar concept that was successfully tested in Arabidopsis (Wijnker et al. 2014). However, this approach involves the use of genetically modified lines with suppressed recombination and the production of doubled-haploid lines by in vitro culture, two methods that are well established in a limited number of crops. Additionally, Reverse Breeding was proposed to generate homozygous parents to obtain a hybrid identical to one already existing, while our goal is the generation of individuals with a similar genetic background but with different characteristics with respect to a top performing clonally-reproducing cultivar.

This strategy is particularly suitable for clonally reproducing species, fruit trees, berries, and ornamentals, among others. Key elements of their biology are the perennial habit of most species and a long intergeneration period, often accompanied by a juvenile period, which requires as few generations as possible to obtain a new cultivar. Our model, peach, has certain advantages: a relatively short intergeneration period (3-4 years), self-compatibility, and a compact, diploid and sequenced genome (250 Mbp; *x*=8) (Verde et al. 2013; Arús et al. 2012), with a low recombination rate (1.2-1.5 crossovers/chromosome and meiosis) and a high level of homozygosity in the commercial cultivars (Micheletti et al. 2015; da Silva Linge et al. 2018). In this paper we develop the basic scheme for Resynthesis, and its genetic foundation, and provide experimental data from a trial using the peach ‘Sweet Dream’ (SD) cultivar. Due to the length of time required for proof of concept, here we provide only data from the marker selection of SD selfed progeny to evaluate the progress made and the feasibility of the approach.

## MATERIALS AND METHODS

### Breeding scheme

#### Resynthesis A

This method consists of selecting two highly homozygous and complementary individuals in the selfed offspring of the original variety that, when crossed, reproduce most of its heterozygous make-up along with additional homozygous regions (Figure 1A). This approach has the advantage that the two selected parents, being highly homozygous, can be used for the introgression of new characters from other sources. The disadvantage is that obtaining these two complementary lines may typically require more than one generation of selfing plus an additional generation to obtain the resynthesized line (or set of lines) and evaluate its behavior.

**Figure 1.**
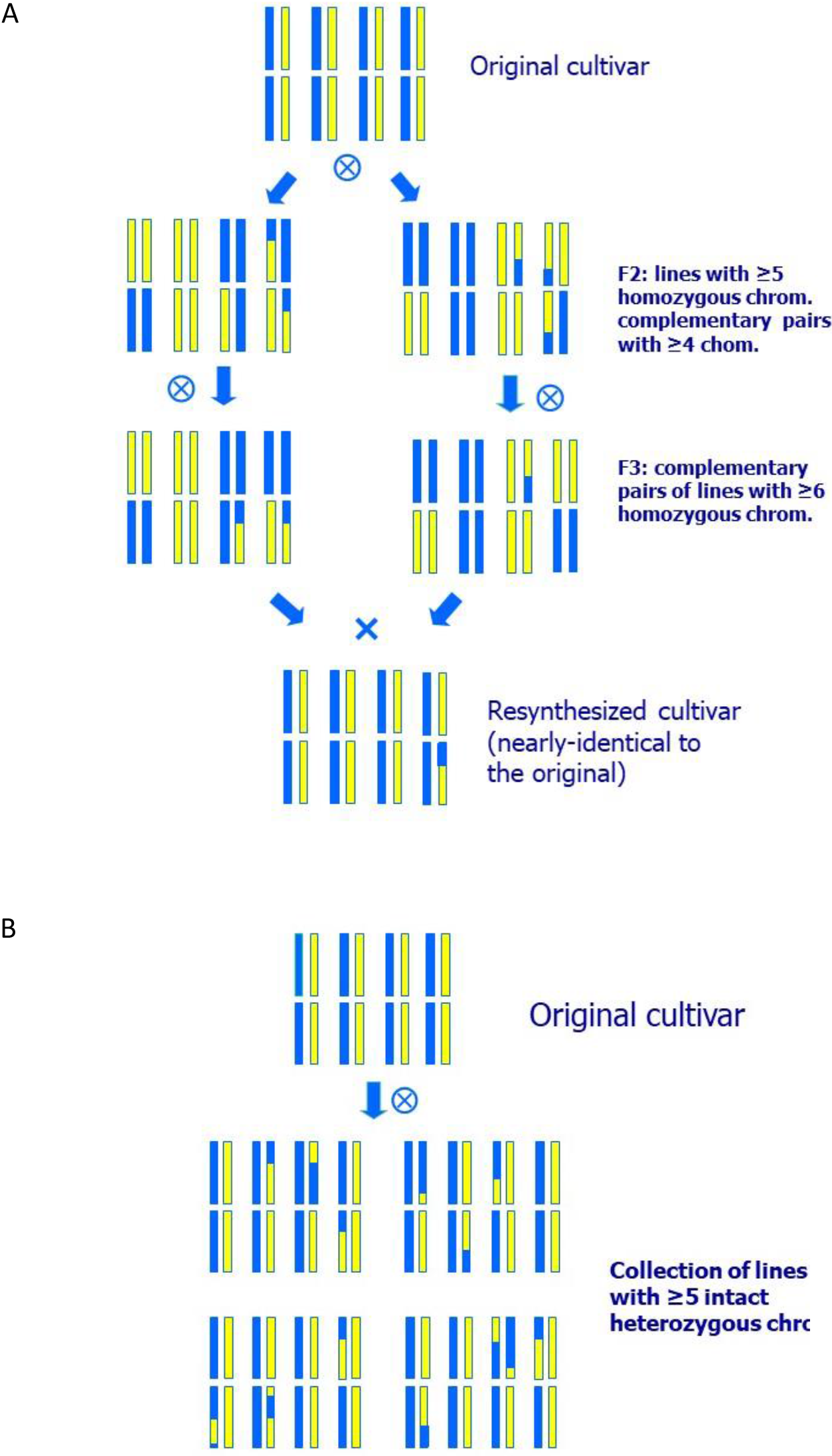
Breeding schemes for A) Resynthesis A, where two nearly homozygous and complementary lines are selected to obtain an individual highly similar to the original cultivar; B) Resynthesis B: Generation of a collection of individuals nearly identical to a peach cultivar, but with a small number of chromosomal segments in homozygosis. All individuals of the collection presented have five heterozygous chromosomes as in the original cultivar and only three homozygous fragments each.

#### Resynthesis B

Using the same F_2_ population as for Resynthesis A, it is possible to select plants with a similar composition to the original cultivar (Figure 1B). For this, markers are used to select individuals with most chromosomes the same as the original variety and others with more homozygous regions, so that the new homozygous regions are a low proportion of the genome (5-20% of the heterozygous genome in homozygosity). The objective is to evaluate the behavior of the selected plants in the field, and their value as potential new varieties. Additionally, such individuals may constitute an interesting resource for genetic studies, consisting of plants containing different homozygous regions in the otherwise partly heterozygous background of the original variety.

### Genetic basis of the Resynthesis process

The proportions of different genotypes expected in an F2 progeny were calculated for peach with *x*=8 chromosomes, initially assuming one crossover per chromosome. In this case, the probability of having one given chromosome pair in homozygosis is 1/8, so the probability of the eight chromosome pairs being homozygous is (1/8)^8^≈6.0×10^-8^, or 1.2×10^-3^ for five or more homozygous chromosome pairs (Table 1; Figure 1) calculated from a binomial distribution with p=1/8. The probabilities are the same for finding individuals entirely heterozygous like the original cultivar, for one or more chromosomes.

**Table 1.** Approximate probabilities of finding a progeny homozygous (or heterozygous) for various intact chromosomes (cs) in the selfed progeny of peach (*x*=8).

|  | Peach |  |  |  |
| --- | --- | --- | --- | --- |
|  | Fully heterozygous parent; 1 co/cs<sup>1</sup> | 50% of the genome identical by descent<sup>2</sup> |  |  |
|  |  | 1 co/cs | 1.2 co/cs | 1.5 co/cs |
| All cs in homozygosis | $6.0 \times 10^{-8}$ | $3.9 \times 10^{-5}$ | $1.3 \times 10^{-5}$ | $2.1 \times 10^{-6}$ |
| All cs in heterozygosis | $6.0 \times 10^{-8}$ | $3.9 \times 10^{-5}$ | $1.3 \times 10^{-5}$ | $2.1 \times 10^{-6}$ |
| Six or more homozygous cs | $8.5 \times 10^{-5}$ | $8.0 \times 10^{-3}$ | $3.8 \times 10^{-3}$ | $1.1 \times 10^{-3}$ |
| Five or more homozygous cs | $1.2 \times 10^{-3}$ | $4.5 \times 10^{-2}$ | $2.5 \times 10^{-2}$ | $9.4 \times 10^{-3}$ |
| Four or more homozygous cs | $1.1 \times 10^{-2}$ | 0.16 | 0.11 | $5.2 \times 10^{-2}$ |

<sup>1</sup>co/cs = crossovers/chromosome; <sup>2</sup> Estimated assuming that half of each chromosome is heterozygous and the other half homozygous

In the case of peach, there is an additional advantage: recent results have shown that vast regions of the chromosomes of the modern cultivars (sometimes accounting for 40-60% of the genome) are monomorphic (Serra et al. 2017, Donoso et al. 2015), identity by descent (IBD) being the most logical cause due to a high level of co-ancestry (Micheletti et al. 2015) of peach commercial cultivars. This determines that some of the crossovers produced at the homozygous DNA fragments will not be effective. As an approximation, if we consider that an individual with 50% IBD has a continuous fragment of half of the length of each chromosome in homozygosis, the probability of a fully homozygous chromosome with a single recombination event increases to 9/32, and the probabilities of having eight or five or more chromosome pairs fully homozygous can be estimated as 3.9×10^-5^ and 8.0×10^-3^, respectively (Table 1). For a finer estimation including recombination rates, we know that in peach the number of crossovers per chromosome (co/cs) per meiosis ranges between 1 and 2, and an overall estimation of 1.2-1.5 is reasonable (Arús et al. 2005; da Silva Linge et al. 2018). We have calculated approximate probabilities of finding a given number of chromosomes in homozygosis (or heterozygosis) considering 1.2 and 1.5 co/cs: 1.2 (80% of the meiosis with a single co/cs and 20% with two) and 1.5 (50% with one co/cs and 50% with two), and 50% IBD (half a chromosome in homozygosis and half in heterozygosis). These calculations indicate that the expected probabilities are compatible with finding some of the desired genotypes in populations of several hundred plants (Table 1).

Considering as an example a peach F2 progeny of N=2,000, with 1.2 co/cs and 50% homozygosity, an average of nine plants with six or more homozygous chromosome pairs would be expected, or 50 plants with five or more homozygous chromosomes. In this case, the objective would be to select 1-5 pairs of plants with four or more chromosomes in homozygosis and complementary to each other. This is achievable because the number of individuals needed to find two complementary lines from a set of lines with four homozygous chromosomes is n=10 with a 0.95 probability. This is calculated with the formula n(n-1)/2=log0.05/log[(2^x^-1)/2^x^], where n is the number of individuals with x (four in this case) homozygous chromosomes. And approximately one in four of the possible pairs of these 50 plants with five homozygous chromosomes would have four homozygous chromosomes in common. It is also important to be sure that the three non-homozygous chromosomes of each pair of lines allow construction of two complementary lines without homozygous fragments of the same haplotype in both lines. By selfing these individuals, sets of genotypes could be obtained that are homozygous for six or more chromosomes and complementary in a sample of N=1,000. From these we would expect to obtain at least one pair of lines that, crossed, would produce a plant ≥ 95% genetically identical to SD with >10% probability.

We could use the same F2 population of N=2,000 to test the Resynthesis B approach by selecting for heterozygous individuals with chromosomal compositions identical to SD, with the same probabilities as those calculated for finding a homozygous genotype (Table 1). A set of 50 plants on average with five or more chromosomes identical to SD can be selected. This set would contain plants having a few homozygous fragments in an essentially heterozygous background allowing genetic analysis of the segregation in these regions by comparison with the original cultivar. Plants of this set can be selected to have the ensemble of homozygous fragments covering ideally the whole genome for each of the two haplotypes, a situation somewhat similar to a near-isogenic line (NIL) collection in a heterozygous background. Some of these lines could be advanced to new cultivars.

### Case study: Resynthesis of cultivar ‘Sweet Dream’ (SD)

#### Plant materials

Fruit from a single stand commercial plantation of SD were collected at maturity in a commercial orchard of Els Alamús (Lleida) in July of 2015, 2016 and 2017 (4,300 fruit in total). Seeds were extracted, cleaned and kept in vacuum bags until November, when they were stratified at 4° C for three months, after which they were germinated in trays in the greenhouse. Germinated plants were transplanted to the field of the IRTA Experimental Station of Gimenells, and grown following standard agricultural practices.

#### Resequencing ‘Sweet Dream’

Young leaves were collected in the spring of 2015 and DNA extracted for resequencing using DNeasy^®^ Plant Mini Kit (Qiagen) following the manufacturer’s instructions. A paired-end sequencing library with an average insert size of 315 bp was prepared following the manufacturer’s instructions (Illumina). The library was sequenced at CNAG (Centre Nacional d’Anàlisi Genòmica, Barcelona) using the Illumina HiSeq2500 platform at a paired-end mode with 2×250 cycles. Adapter removal and read trimming was done using Trimmomatic v0.36 (Bolger et al., 2014). FastQC (version 0.10.1; https://www.bioinformatics.babraham.ac.uk/projects/fastqc/) was used for quality control of the reads before and after adapter removal and trimming. With a good quality dataset, reads were mapped onto the peach genome downloaded from GDR (ftp://ftp.bioinfo.wsu.edu/species/Prunus_persica/Prunus_persica-genome.v2.0.a1/assembly/Prunus_persica_v2.0.a1_scaffolds.fasta.gz) using BWA-MEM (v0.7.5a) (Li and Durbin., 2009). Alignment files were sorted and reads that were uniquely mapped to the genome were retained for variant calling using Samtools v1.3.0 (Li 2011), taking into account reads with minimum base quality of 10 and minimum mapping quality of 20. From the list of variants, we removed INDELs, sites where the quality of the SNP and the genotype quality were lower than 30, sites with depth lower than 12, sites with more than one alternative allele and, finally, sites where there was a ratio imbalance >0.3 between forward and reverse reads of the same allele. In order to keep sites with high genotype qualities we removed those sites where the heterozygous SNPs had an allele frequency (AF) ratio outside the 0.4 – 0.6 range and the homozygous SNPs had an AF ratio lower than 0.8.

### Fluidigm SNP array development, genotyping, mapping and selection in the SD F2 population

To develop a 96 SNP array to genotype the SD F2 seedlings, SNPs covering the whole genome were selected. We chose 87 known to be heterozygous in SD and previously validated in other peach materials from the peach 9k array (Verde et al. 2012; Micheletti et al. 2015). For genomic regions not covered by these markers, we identified nine more heterozygous SNPs in SD from the resequencing data. The complete SNP list with their main characteristics is given in Tables S1 and S2.

Fresh leaves from SD and from its F2 progeny were collected directly in 96 well plates. DNA was extracted following the Doyle & Doyle (1990) protocol. Genotyping was carried out with the KASPar SNP Genotyping System (KBiosciences, Herts, UK) adapted to a Biomark™ HD platform (Fluidigm) and using the universal KASPar MasterMix (LGC, England) following the instructions recommended by the supplier. KASPar assay primers were designed with Primer Picker (KBiosciences) (Table 1S), and SNP data were analysed with the Fluidigm SNP Genotyping Analysis software (Fluidigm).

Markers with less than 90% of the data available were discarded. An initial marker order was established according to their physical position and SNPs located on the same chromosome were placed in the same phase. Missing data were imputed in chromosome fragments that were delimited by markers with the same genotype. Imputations were limited to 1.6% of the data available. Markers were then mapped with Kosambi’s genetic distance using MapMaker 3.0 (Lander et al. 1987), and MapChart 2.1 (Voorrips 2002) was used to draw the resulting linkage map.

Only mapped markers were used for the selection for Resynthesis A and B. For each plant, we calculated the frequency of heterozygous loci and ordered the plants according to this parameter. For selection of plants with complementary genotypes, we identified first the plants with heterozygosity values below 0.25 and looked for possible complementary pairs by selecting those with higher distance values among those with heterozygosity values below 0.50. Distance was calculated for each locus as 0 for markers with the same genotype, 0.5 for heterozygous vs. homozygous comparisons and 1 for the two alternative homozygotes and summed over all loci. Only pairs of genotypes with high distances that were totally compatible, i.e. where the same allele was not fixed at any given locus for both parents, were accepted as possible parents for the next generation.

## RESULTS

From the approximately 4,300 seeds stratified, 416 (135 in 2015, 136 in 2016 and 145 in 2017), germinated and produced viable plants. This is a low germination rate (~10%), considering that we used the standard methods of IRTA’s peach breeding program, where the average germination rate is 20-30%, although it is known that certain parents and crosses produce very poor yields, and that there is an extremely wide range of success (0-90%), depending on the cross. One of the reasons we selected SD was because, being a mid-season producing cultivar, no need for ‘in vitro’ embryo rescue was expected, as occurs in early varieties. We attribute this low rate of germination to the characteristics of the cultivar, requiring specific adaptation of the germination protocol to improve results.

Illumina sequencing of SD DNA generated 91,441,354 reads with a total output of 22.8Gbp. After quality filtering we were left with 81,866,684. Of these, 66,950,399 (81.8%) were mapped onto the peach genome, covering 94% of its length with at least five reads. The resulting average depth was determined to be 36X. In total, we detected 94,451 heterozygous SNPs, with chromosomes Pp02 having the highest and Pp08 the lowest number, and with an average of 11,863 heterozygous SNPs per chromosome (Table S3). This corresponds to an average SNP density of 44.7 SNPs/Mbp, with Pp02 and Pp01 having the highest (72.6) and lowest (14.1) density, respectively. Looking at the distribution of heterozygous SNPs along the chromosomes, they are concentrated in certain parts of the genome, leaving large of interspersed homozygous regions, particularly in chromosomes 1, 3, 6 and 8 (Figure S1). Considering that regions below 5% of the average of SNPs/Mbp (21 SNP/Mbp; see Table 1) are identical by descent, we calculate that the genome of SD is 23% IBD, in agreement with the high level of inbreeding of peach cultivars (Micheletti et al. 2015).

From the 96 SNPs used for analysis, 64 segregated and were mapped and of the remaining 32, 16 were monomorphic and 16 had no call or an unclear segregation pattern. Of the 64 mapped SNPs, 62 were codominant with the three expected genotype classes and two (SNP_IGA_671806 and SNP_IGA_682254) were dominant, with only two genotypes. A map was constructed that detected all eight linkage groups corresponding to the expected eight *Prunus* chromosomes. The order of the markers within the linkage groups was identical to their physical order in the chromosomes (see Table S1). The map covered a total distance of 422.5 cM, ranging from 36.0-63.7 cM/chromosome and the markers studied covered 71% of the total physical length of the peach genome. Two of the main regions uncovered by the map, end of chromosome 1 (21.3 Mbp) and end of chromosome 7 (4.2 Mbp), were predominantly homozygous (Figure S1) in SD.

For Resynthesis A, we selected four pairs of complementary lines with average heterozygosity of the two parents ≤0.27 (Figure 2B). These lines had on average 3.3 chromosomes in homozygosity ranging from 0 to 8, and the average number of estimated recombinations per genome was 4.8 (0.6 co/cs), ranging from 2 to 8. Recovering entire chromosomes without any recombinations would be the ideal situation, because the presence of one or more recombination events almost necessarily implies that there will be a more or less large homozygous region in the resynthesized genotype, as crossovers rarely occur at the same genomic site in two meiosis.

**Figure 2.**
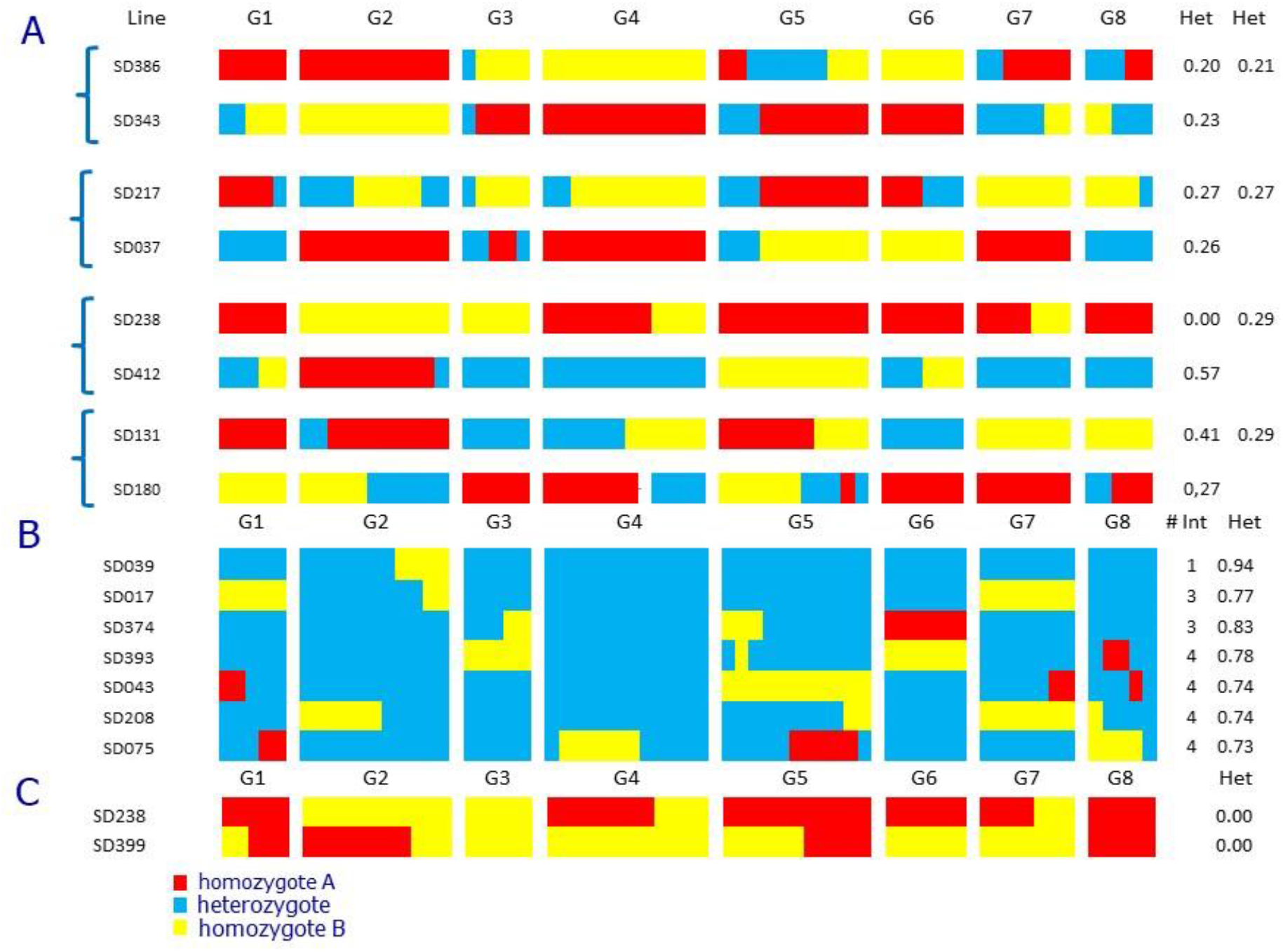
Chromosome diagram of ‘Sweet Dream’ F_2_ selected individuals. G1-G8 peach linkage groups. Het: proportion of heterozygous loci. # Int: introgression number A) Pairs of nearly-homozygous lines selected for Resynthesis A; with high probability their offspring will produce individuals very close to ‘Sweet Dream’. B) Selected individuals for Resynthesis B, nearly identical to ‘Sweet Dream’. C) Fully homozygous F2 plants.

For Resynthesis B, we found seven individuals with >73% heterozygosity (range 73-94%) that contained the full genome of SD and a few DNA fragments that were fixed (Figure 2A). One of them contained a single homozygous fragment, two contained three and the remaining four plants, four.

When looking for individuals with high levels of homozygosity, we found two that were completely homozygous for all markers studied (Figure 2C). One of them (SDF2-238) was selected as one of the parents of the complementary lines. Both plants had two recombination events, meaning that in the regions where recombination occurred there would be a small heterozygous DNA fragment, the rest of the genome being fully homozygous.

## DISCUSSION

A major use of molecular marker analysis is to predict the phenotype based on genotypic data, which has important applications in plant breeding. One of the critical advantages of this strategy is that, once the marker-phenotype association has been established, it is possible to genotype large progenies at the seedling stage, and only a few selected individuals need to be grown and phenotyped. In the case of Resynthesis, the target genotype is known and the objective is to obtain, in the shortest possible timeframe, individuals having a similar genetic makeup, some maintaining most of its good phenotypic qualities, but with commercially relevant differences. These differences would be initially provided by the inter-allelic variability of the same cultivar, but in addition, Resynthesis A would give access to alleles of a broader germplasm pool by introgressing them in the inbred parents that would again generate the target genotype.

In spite of the relatively low number of seedlings analyzed, we found four pairs of complementary lines which were suitable for a second generation of Resynthesis A. We expect to perform the first crosses with these lines in the spring of 2021 to obtain populations that will be genotyped and integrated in our breeding program for the following years, to select for individuals similar to SD, with specific characters incorporated. These four pairs of parents will also be selfed, allowing for additionally fixed (5-6 chromosomes in homozygosity) and complementary individuals to be selected.

We found seven individuals nearly identical to SD (Resynthesis B), which gives us an opportunity to look for the effects of specific regions of the genome on key agronomic characters, although to a limited extent (six genomic regions in total covering chromosome 6 and parts of chromosomes 1, 5, 7 and 8). More individuals will be produced with appropriate compositions from the crosses between the complementary pairs of Resynthesis A that, once genotyped, will allow for additional genome coverage. It would not be surprising if some of these individuals have characteristics that may already make them commercially interesting for selection as new cultivars.

The two completely homozygous lines were an unexpected result, showing that these individuals can be recovered with a limited number of offspring. One of them was selected as one of the parents for Resynthesis A, but beyond that, these lines are of biological interest as a relatively rare event. The self-pollinating nature of peach provides the opportunity to produce inbreds, and certain traditional cultivars of Spain are highly homozygous (Li et al. 2013; Micheletti et al. 2015) as a consequence of their seed multiplication before grafting techniques became sufficiently well established. Growing a number of progeny from self-compatible cultivars of other fruit trees (almond, cherry, apricot) could produce inbreds that may be of use for genetic analysis, including ‘de novo’ sequencing.

Massive linkage disequilibrium is a favorable circumstance for Resynthesis and species with low chromosome numbers and low recombination rates, usually associated with small genomes (Mercier et al. 2005), are good candidates for a fast Resynthesis process. To measure haplotype integrity after meiosis, we propose a recombination index Ri = n(1+CpC), where n is the number of chromosomes and CpC the average number of crossovers per chromosome that can be inferred from linkage maps available. Species with low Ri would require less individuals and/or less generations to obtain suitable pairs of inbreds for Resynthesis. Peach, for example, with eight chromosomes and 1.3 co/cs (da Silva Linge 2018) would have a Ri =8+8×1.3=18.4. Almond (*x*=8, 1.0 co/cs) Ri=16.0 (Sánchez-Pérez at al. 2007), raspberry (*x*=7, 1.3 co/cs) Ri=16.1 (Ward et al. 2013) or citrus (*x*=9, 1.7 co/cs) Ri= 24.3 (Huang et al. 2018) are examples of favorable organisms for Resynthesis, whereas apple (*x*=17 1.5 co/cs) Ri=42.5 (di Pierro et al. 2016) or grape (*x*=19; 1.5 co/cs) Ri= 47.5 (Tello et al. 2019), would require greater population sizes to obtain sufficiently close individuals. The capability of each species to produce large numbers of seedlings per generation at affordable costs is also important: in this respect grapes and apple are more amenable to Resynthesis than peach. Hybrid populations between pairs of complementary lines could be obtained using partly homozygous lines from early steps of selfing (i.e. from F_2_ selected couples), and plants with valuable phenotypes selected at this stage. Selfing generations to fix specific lines could be done in parallel. Backcrossing of selected lines to obtain parents with genes from other materials (disease resistance or others), could also be started at early stages.

It is important that the genotype to be resynthesized is self-compatible as the complementary lines are obtained from successive selfing generations. For self-incompatible genotypes, a possible way to circumvent the problem is crossing first with a self-compatible genotype, select the self-compatible individual in this progeny with the least number of recombination events in the original cultivar gamete, and start the selection from the BC1 between the hybrid line and the target genotype. Then only self-compatible lines having as much as possible of their genome fixed for one of the alleles of the target genotype (looking for pairs of complementary lines following a model similar to Resynthesis A) or that are highly similar to its genetic makeup (Resynthesis B), should be selected.

Another aspect to consider is the existence of lethal alleles in heterozygosis in the original cultivar. This cannot be discarded in highly heterozygous genomes of typically outcrossing species, such as those of almond and grape, especially in old cultivars of unknown origin. In this case, one of the two complementary lines must be kept in heterozygosis for the smallest possible genomic region around the gene causing the lethal effect, and selection should be for heterozygous individuals in the final cross between the two nearly-homozygous lines in the last step of Resynthesis A. Knowing whether a certain cultivar has lethal or deleterious alleles in heterozygosis could be tested by constructing a linkage map with markers in the F2 and identifying genomic regions with highly distorted segregation frequencies. No regions of this type have been detected in our F2 population of SD.

Resynthesis could be expanded to other crops, particularly those that are seed propagated and where F1 hybrids are the usual cultivar type. In this case, there is an alternative approach to the usual generation of inbred lines and selection of the best performing F1 hybrids produced between them: whenever the evaluation of single genotypes (individually or as a set of clonally propagated plants) from segregating progenies can be done with sufficient accuracy, highly performing individuals could be selected and their genotype A resynthesized, following a strategy that somewhat mimics the clonally-propagated breeding scheme.

## ACKNOWLEDGMENTS

This research was supported in part by grants from: the Spanish Ministry of Economy and Competitiveness (MINECO/FEDER projects AGL2012-40228-C02-01 and RTA2015-00050-00-00), the project RIS3CAT (COTPA-FRUIT3CAT) financed by the European Regional Development Fund through the FEDER frame of Catalonia 2014-2020, the Severo Ochoa Program for Centres of Excellence in R&D 201-2019 (SEV-2015-0533) and CERCA Programme-Generalitat de Catalunya.

## REFERENCES

Arús P, Yamamoto T, Dirlewanger E, Abbott AG (2005) Synteny in the Rosaceae. Plant Breeding Reviews, 27. Jules Janick (ed.) pp. 175–211.

Arús P, Verde I, Sosinski B, Zhebentyayeva T, Abbott AG (2012) The peach genome. Tree Genet Genomes 8:531–547

Bolger AM, Lohse M, Usadel B (2014) Trimmomatic: a flexible trimmer for Illumina sequence data. Bioinformatics 30:2114–2120

da Silva Linge C, Antanaviciute L, Abdelghafar A, Arús P, Bassi D, Rossini L, Flickin F, Gasic K (2018) High-density multi-population consensus genetic linkage map for peach. PLoS ONE 13(11): e0207724.

Di Pierro EA, Gianfranceschi L, Di Guardo M, Koehorst-van Putten HJ, Kruisselbrink JW, Longhi S, Troggio M, Bianco L, Muranty H, Pagliarani G, Tartarini S, Letschka T, Lozano Luis L, Garkava-Gustavsson L, Micheletti D, Bink MC, Voorrips RE, Aziz E, Velasco R, Laurens F, van de Weg WE (2016) A high-density, multi-parental SNP genetic map on apple validates a new mapping approach for outcrossing species. Hortic Res 3:16057

Dirks R, van Dun K, de Snoo CB, van den Berg M, Lelivelt CLC, Voermans W, Woudenberg L, de Wit JPC, Reinink K, Schut JW, van der Zeeuw E, Vogelaar A, Freymark G, Gutteling EW, Keppel MN, van Drongelen P, Kieny M, Ellul P, Touraev A, Ma H, de Jong H and Wijnker E (2009) Reverse breeding: a novel breeding approach based on engineered meiosis. Plant Biotechnol J 7: 837–845

Donoso JM, Picañol R, Eduardo I, BatlIe I, Howad W, Aranzana MJ, Arús P (2015) High-density mapping suggests a cytoplasmic male sterility system with two restorer factors in almond x peach progenies. Hortic Res 2, 15016

Doyle JJ, Doyle JL (1990) Isolation of plant DNA from fresh tissue. Focus 12:13–15

Huang M, Roose ML, Yu Q, Du D, Yu Y, Zhang Y, Deng Z, Stover E, Gmitter FG (2018) Construction of High-Density Genetic Maps and Detection of QTLs Associated With Huanglongbing Tolerance in Citrus. Front Plant Sci 9:1694.

Koumproglou R, Wilkes TM, Townson P, Wang XY, Beynon J, Pooni HS, Newbury HJ, Kearsey MJ (2002) STAIRS: a new genetic resource for functional genomic studies of Arabidopsis. Plant J 31(3):355–364

Lander, E S, Green, P, Abrahamson, J, Barlow, A, Daly, M J, Lincoln, S E and Newburg, L (1987) MAPMAKER: An interactive computer package for constructing primary genetic linkage maps of experimental and natural populations. Genomics 1: 174–181

Li H (2011) A statistical framework for SNP calling, mutation discovery, association mapping and population genetical parameter estimation from sequencing data. Bioinformatics, 27:2987–2993

Li H, Durbin R (2009) Fast and accurate short read alignment with Burrows-Wheeler transform. Bioinformatics 25:1754–1760

Li XW, Meng XQ, Jia HJ, Yu ML, Ma RJ, Wang LR, Cao K, Shen ZJ, Niu L, Tian JB, Chen MJ, Xie M, Arús P, Gao ZS, Aranzana MJ (2013) Peach genetic resources: diversity, population structure and linkage disequilibrium. BMC Genetics 14:84

Mercier R., Mezard C., Jenczewski E., Macaisne N., Grelon M. (2015) The Molecular Biology of Meiosis in Plants, in: S. S. Merchant (Ed.), Ann Rev Plant Biol 66: 297–327.

Micheletti D, Dettori MT, Micali S, Aramini V, Pacheco I, Da Silva Linge C, Foschi S, Banchi E, Barreneche T, Quilot-Turion B, Lambert P, Pascal T, Iglesias I, Carbó J, Wang L-R, Ma R-J, Li X-W, Gao Z-S, Nazzicari N, Troggio M, Bassi D, Rossini L, Verde I, Laurens F, Arús P, Aranzana MJ (2015) Whole-genome analysis of diversity and SNP-major gene association in peach germplasm. PLoS ONE 10.

Sánchez-Pérez R, Howad W, Dicenta F, Arús P, Martínez-Gómez P (2007) Mapping major genes and quantitative trait loci controlling agronomic traits in almond. Plant Breed 125:1–9

Serra O, Giné-Bordonaba J, Eduardo I, Bonany J, Echeverría G, Larrigaudière G, Arús P (2017) Genetic analysis of the slow melting flesh character in peach. Tree Genet Genomes 13:77

Tello J, Roux C, Chouiki H, Laucou V, Sarah G, Weber A, Santoni S, Flutre T, Pons T, This P, Péros JP, Doligez A (2019) A novel high-density grapevine (Vitis vinifera L.) integrated linkage map using GBS in a half-diallel population. Theor Appl Genet 132:2237–2252

Verde I, Bassil N, Scalabrin S, Gilmore B, Lawley CT, Gasic K, Micheletti D, Rosyara UR, Cattonaro F, Vendramin E, Main D, Aramini V, Blas AL, Mockler TC, Bryant DW, Wilhelm L, Troggio M, Sosinski B, Aranzana MJ, Arús P, Iezzoni A, Morgante M, Peace C (2012) Development and evaluation of a 9k SNP array for peach by internationally coordinated snp detection and validation in breeding germplasm. PLoS ONE 7:

Verde I, Abbott AG, Scalabrin S, Jung S, Shu S, Marroni F, Zhebentyayeva T, Dettori MT, Grimwood J, Cattonaro F, Zuccolo A, Rossini L, Jenkins J, Vendramin E, Meisel LA, Decroocq V, Sosinski B, Prochnik S, Mitros T, Policriti A, Cipriani G, Dondini L, Ficklin S, MGoodstein D, Xuan P, Del Fabbro C, Aramini V, Copetti D, Gonzalez S, Horner DS, Falchi R, Lucas S, Mica E, Maldonado J, Lazzari B, Bielenberg D, Pirona R, Miculan M, Barakat A, Testolin R, Stella A, Tartarini S, Tonutti P, Arús P, Orellana A, Wells C, Main D, Vizzotto G, Silva H, Salamini F, Schmutz J, Morgante M, Rokhsar DS (2013) The high-quality draft genome of peach (*Prunus persica*) identifies unique patterns of genetic diversity, domestication and genome evolution. Nature Genet 45(5):487–494

Voorrips RE (2002) MapChart: Software for the graphical presentation of linkage maps and QTLs. J Hered 93: 77–78.

Ward JA, Bhangoo J, Fernandez-Fernandez F, Moore P, Swanson JD, Viola R, Velasco R, Bassil N, Weber CA, Sargent DJ (2013) Saturated linkage map construction in Rubus idaeus using genotyping by sequencing and genome-independent imputation BMC Genomics 14:2

Wijnker E, Deurhof L, van de Belt J, de Snoo CB, Blankestijn H, Becker F, Ravi M, Chan SWL, van Dun K, Lelivelt CLC, de Jong H, Dirks R, Keurentjes JJB (2014) Hybrid recreation by reverse breeding in Arabidopsis thaliana. Nature Protocols 9:761–772.

