## Supplemental Materials for "Resynthesis: Marker-based partial reconstruction of elite genotypes in clonally-reproducing plant species"

**Table S1**. Description of the peach SNP markers included in the Fluidigm chip used for the construction of the 'Sweet dream' (SD) map

| SNP name | Abbreviation (Map) | Chromosome | Physical position (bp) | SNP type | SD map | cM[1] |
|---|---|---|---|---|---|---|
| SNP_Pp01-8080 | na[2] | Pp01 | 8080 | A/C | monomorph | um[3] |
| SNP_IGA_2651 | na | Pp01 | 883576 | T/C | No call | um |
| SNP_IGA_17419 | SI17419 | Pp01 | 6000280 | T/C | yes | 0.0 |
| SNP_IGA_19514 | SI19514 | Pp01 | 6872054 | T/C | yes | 4.5 |
| SNP_IGA_31646 | na | Pp01 | 12187249 | T/C | monomorph | um |
| SNP_Pp01-15435423 | na | Pp01 | 15435423 | A/G | monomorph | um |
| SNP_IGA_67137 | SI67137 | Pp01 | 20219128 | T/G | yes | 25.4 |
| SNP_IGA_79455 | SI79455 | Pp01 | 23529157 | T/C | yes | 30.1 |
| SNP_IGA_86968 | SI86968 | Pp01 | 26640558 | T/G | yes | 36 |
| SNP_IGA_103771 | na | Pp01 | 33744976 | A/G | monomorph | um |
| SNP_IGA_120784 | na | Pp01 | 41413142 | T/C | No call | um |
| SNP_IGA_125219 | na | Pp01 | 46984218 | T/C | no | um |
| SNP_IGA_123719 | na | Pp01 | 47532400 | A/G | No call | um |
| SNP_IGA_238135 | na | Pp02 | 241474 | T/C | No call | um |
| SNP_IGA_136625 | SI136625 | Pp02 | 2002076 | A/G | yes | 0.0 |
| SNP_IGA_148760 | SI148760 | Pp02 | 3109681 | A/G | yes | 3.3 |
| SNP_IGA_195134 | SI195134 | Pp02 | 6545323 | A/G | yes | 9.6 |
| SNP_IGA_218801 | SI218801 | Pp02 | 9053696 | T/C | yes | 13.1 |
| SNP_IGA_242118 | SI242118 | Pp02 | 13717350 | A/G | yes | 17.7 |
| SNP_IGA_252957 | SI252957 | Pp02 | 15849694 | A/G | yes | 23.6 |
| SNP_IGA_258854 | SI258854 | Pp02 | 17219077 | A/G | yes | 26.8 |
| SNP_IGA_266063 | SI266063 | Pp02 | 19744284 | T/G | yes | 34.3 |
| SNP_IGA_275502 | SI275502 | Pp02 | 22301505 | T/C | yes | 41.1 |
| SNP_IGA_283325 | SI283325 | Pp02 | 25849495 | T/C | yes | 52.5 |
| SNP_IGA_285058 | SI285058 | Pp02 | 26895362 | A/G | yes | 58.8 |
| SNP_IGA_898654 | SI898654 | Pp03 | 742389 | A/G | yes | 0.0 |
| SNP_IGA_299796 | na | Pp03 | 4209014 | A/C | monomorph | um |
| SNP_Pp03-3917979 | na | Pp03 | 3917979 | T/C | No call | um |
| SNP_IGA_322698 | na | Pp03 | 9911321 | A/G | monomorph | um |
| SNP_IGA_428056 | na | Pp03 | 11876389 | T/C | monomorph | um |
| SNP_IGA_894039 | na | Pp03 | 13664157 | T/C | No call | um |
| SNP_IGA_325850 | SI325850 | Pp03 | 15520683 | A/C | yes | 27.6 |
| SNP_IGA_344612 | SI344612 | Pp03 | 19098840 | A/G | yes | 35.9 |
| SNP_IGA_349233 | SI349233 | Pp03 | 20300608 | T/C | yes | 43.9 |
| SNP_IGA_356179 | SI356179 | Pp03 | 22294942 | A/G | yes | 53.4 |

[1]cM= genetic distance in centimorgans
[2]na=not assigned
[3]um=unmapped

**Table S1** (continued)

| SNP name | Abbreviation (Map) | Chromosome | Physical position (bp) | SNP type | SD map | cM |
|---|---|---|---|---|---|---|
| SNP_IGA_368926 | SI368926 | Pp04 | 407324 | T/C | yes | 0.0 |
| SNP_IGA_379856 | SI379856 | Pp04 | 1477791 | T/C | yes | 3.5 |
| SNP_IGA_389204 | SI389204 | Pp04 | 4843247 | A/G | yes | 15 |
| SNP_IGA_395253 | SI395253 | Pp04 | 6202308 | T/C | yes | 21.1 |
| SNP_IGA_398213 | SI398213 | Pp04 | 6895567 | A/G | yes | 25.5 |
| SNP_IGA_403741 | SI403741 | Pp04 | 9106604 | A/G | yes | 33.2 |
| SNP_IGA_409453 | SI409453 | Pp04 | 10396616 | T/C | yes | 37.7 |
| SNP_IGA_420819 | na | Pp04 | 13949755 | A/C | monomorph | um |
| SNP_Pp04-13754607 | na | Pp04 | 13754607 | A/G | monomorph | um |
| SNP_IGA_440116 | SI440116 | Pp04 | 16084694 | T/G | yes | 52 |
| SNP_IGA_454568 | SI454568 | Pp04 | 18760831 | A/G | yes | 57.3 |
| SNP_IGA_480804 | SI480804 | Pp04 | 20068418 | A/G | yes | 58.2 |
| SNP_IGA_513502 | SI513502 | Pp04 | 22616919 | A/G | yes | 59.5 |
| SNP_IGA_528949 | SI528949 | Pp04 | 24187959 | A/G | yes | 63.7 |
| SNP_IGA_544495 | SI544495 | Pp05 | 610569 | T/G | yes | 0.0 |
| SNP_IGA_548037 | SI548037 | Pp05 | 1376475 | T/C | yes | 4.1 |
| SNP_IGA_553456 | SI553456 | Pp05 | 2477309 | A/G | yes | 8.2 |
| SNP_IGA_571548 | SI571548 | Pp05 | 5485331 | A/G | yes | 17.4 |
| SNP_IGA_572589 | SI572589 | Pp05 | 5813029 | A/G | yes | 18.6 |
| SNP_IGA_585810 | SI585810 | Pp05 | 9347092 | T/C | yes | 22.3 |
| SNP_IGA_591439 | SI591439 | Pp05 | 11196070 | A/G | yes | 34.7 |
| SNP_IGA_595126 | SI595126 | Pp05 | 12682824 | T/C | yes | 42.9 |
| SNP_IGA_596393 | SI596393 | Pp05 | 13411012 | T/C | yes | 45.9 |
| SNP_IGA_600493 | SI600493 | Pp05 | 14879113 | A/C | yes | 53.5 |
| SNP_IGA_602605 | SI602605 | Pp05 | 16636368 | T/C | yes | 58.9 |
| SNP_IGA_616458 | na | Pp06 | 1304512 | A/G | No call | um |
| SNP_Pp06-2176846 | na | Pp06 | 2176846 | T/C | monomorph | um |
| SNP_IGA_610889 | na | Pp06 | 3034364 | T/C | No call | um |
| SNP_IGA_627328 | na | Pp06 | 7550350 | T/G | monomorph | um |
| SNP_Pp06-7216093 | na | Pp06 | 7216093 | A/C | monomorph | um |
| SNP_IGA_639062 | na | Pp06 | 11085127 | A/C | monomorph | um |
| SNP_IGA_640395 | SI640395 | Pp06 | 11298491 | T/C | yes | 0.0 |
| SNP_IGA_655825 | SI655825 | Pp06 | 14681941 | T/C | yes | 5.4 |
| SNP_IGA_663057 | SI663057 | Pp06 | 17959563 | A/C | yes | 11 |
| SNP_IGA_671806 | SI671806 | Pp06 | 21011215 | T/C | yes, dominant | 18.5 |
| SNP_IGA_682254 | SI682254 | Pp06 | 24511760 | T/C | yes, dominant | 29.7 |
| SNP_IGA_691624 | SI691624 | Pp06 | 27207318 | A/C | yes | 42.8 |
| SNP_IGA_696341 | SI696341 | Pp06 | 28500652 | T/C | yes | 46.4 |
| SNP_IGA_700653 | SI700653 | Pp06 | 30012439 | A/C | yes | 52 |
| SNP_IGA_726222 | na | Pp07 | 350629 | A/G | No call | um |
| SNP_IGA_717591 | na | Pp07 | 2354039 | T/C | No call | um |
| SNP_IGA_704075 | na | Pp07 | 4239920 | A/C | No call | um |

**Table S1** (continued)

| SNP name | Abbreviation (Map) | Chromosome | Physical position (bp) | SNP type | SD map | cM |
|---|---|---|---|---|---|---|
| SNP_Pp07-5647370 | S0756473 | Pp07 | 5827566 | A/G | yes | 0.0 |
| SNP_IGA_752104 | SI752104 | Pp07 | 8182487 | T/C | yes | 4.5 |
| SNP_IGA_758767 | SI758767 | Pp07 | 10388929 | A/C | yes | 12 |
| SNP_IGA_774557 | SI774557 | Pp07 | 13833942 | A/G | yes | 21.7 |
| SNP_IGA_776826 | SI776826 | Pp07 | 14891881 | A/G | yes | 27 |
| SNP_IGA_781003 | SI781003 | Pp07 | 16422167 | A/G | yes | 38.4 |
| SNP_IGA_784777 | SI784777 | Pp07 | 18244540 | A/G | yes | 46.2 |
| SNP_IGA_794167 | na | Pp08 | 373237 | A/G | Not clear | um |
| SNP_IGA_797492 | na | Pp08 | 1248336 | T/G | monomorph | um |
| SNP_IGA_803699 | na | Pp08 | 2564296 | A/G | Not clear | um |
| SNP_Pp08-3702593 | na | Pp08 | 3702593 | T/C | monomorph | um |
| SNP_IGA_821894 | SI821894 | Pp08 | 5071328 | T/C | yes | 0.0 |
| SNP_IGA_851849 | SI851849 | Pp08 | 10999017 | T/C | yes | 11.1 |
| SNP_IGA_860815 | na | Pp08 | 13540876 | A/G | monomorph | um |
| SNP_Pp08-13670362 | na | Pp08 | 13670362 | T/C | No call | um |
| SNP_IGA_871082 | SI871082 | Pp08 | 16740766 | A/G | yes | 28.3 |
| SNP_IGA_878044 | SI878044 | Pp08 | 18644875 | A/G | yes | 38.6 |
| SNP_IGA_883524 | na | Pp08 | 20833164 | T/G | Not clear | um |
| SNP_IGA_884755 | SI884755 | Pp08 | 21455147 | A/G | yes | 53.5 |

**Table S2**. Sequences of the KASPar assay primers and chromosomal region where the SNPs used in this paper

| SNP name | A1 | A2 | C1 | Sequence |
|---|---|---|---|---|
| SNP_Pp01-8080 | GAAGGTGACCAAGTTCATGCTGTGAAAACACCGTATCTTAACAGTAAGTT | GAAGGTCGGAGTCAACGGATTGAAAACACCGTATCTTAACAGTAAGTG | ATGTTGCGTTCTCAGATATATAGGCCTTT | CTTACCGGTAAGATATGATGTTTTTAGAAAAACAAGTTATTTCCTCTATCAATTACGAGTAAGATGTTGCGTTCTCAGATATATAGGCCTTTTACACTAT[A/C]ACTTACTGTTAAGATACGGTGTTTTCACAAAAATAAGTTATGTAGTCCACGAATTACGGGTAAGACGTGGGGTTCTCATATATCCACGGTTTTACACTAT |
| SNP_IGA_2651 | GAAGGTGACCAAGTTCATGCTCACTCACCACCATGTTGTTATTATTTG | GAAGGTCGGAGTCAACGGATTCCACTCACCACCATGTTGTTATTATTTA | GGGAAAGTGATGGAAGCATTTTGTTTGAA | CATTGAGTATCTCAATTGGTAGTTGCAGATCTTCTTCAACTGCTGAGGGCTCTCATGGGGAAAGTGATGGAAGCATTTTGTTTGAAGAACAAAGAGAAGC[C/T]AAATAATAACAACATGGTGGTGAGTGGAAGAGGAAGCAGTAAAATCAGAGGAGAAGTAGTGTTGATGAAGAAGAATGTAATGGACTTCACTGATATCAGA |
| SNP_IGA_17419 | GAAGGTGACCAAGTTCATGCTGTGCAATGCCCAACAGGTGTC | GAAGGTCGGAGTCAACGGATTCGTGCAATGCCCAACAGGTGTT | GTGATCATTGTGGTGAAAAGAAACATACTA | GCGACAATAGCCACATGAGCTTTATCGATCAGTTGTTCCTTTTTTAAACGCCTCTATTCCTTCTCCTAATCTAGTACCTCGTGCAATGCCCAACAGGTGT[C/T]GATAGTATGTTTCTTTTCACCACAATGATCACACTTTAGTTTGCTTTTTTACAGAAGTCATGCGACAAGGTAGAACGGTAGAAGAGCCAAACCCTATAGG |
| SNP_IGA_19514 | GAAGGTGACCAAGTTCATGCTCATTTTCTGGGGTAGTGGCACC | GAAGGTCGGAGTCAACGGATTAACATTTTCTGGGGTAGTGGCACT | CTGCCTGAGCCGGCGTCCA | CAAGGTATGCACAATGGTATGCAAACCAAAGGCGATACGACTGCGCTTTGAGGCACTCAAACGGCCCTTATGGAGAGAACATTTTCTGGGGTAGTGGCAC[C/T]GGGTGGACGCCGGCTCAGGCAGTGGCGGCATGGGCTTCGGAGAGCAGGTGGTACAATTACTGGTCTAATTCTTGTGCTAGAGGGCAGGAATGTGGGCATT |
| SNP_IGA_31646 | GAAGGTGACCAAGTTCATGCTATTTGACAACAAGAGTGAGTCTGGC | GAAGGTCGGAGTCAACGGATTATATTTGACAACAAGAGTGAGTCTGGT | CATCCGAATTAGCTTCTGGTGGTGA | TTGTTATATCAATTGATGTTTAAAGTACGCAAATTTGATTGATATGTAGTTGATTATTCTGTAAACAGGTGGTGATATTTGACAACAAGAGTGAGTCTGG[C/T]TCACCACCAGAAGCTAATTCGGATGAGGATGAAAGTGGTGCTGCTTTTCTTGTACGAATCTTGAGGTACCTTGAGACACCTCAATATTTGAGAAAAGCTC |
| SNP_Pp01-15435423 | GAAGGTGACCAAGTTCATGCTAACTCGCACAAACGGAAGGATAGA | GAAGGTCGGAGTCAACGGATTCTCGCACAAACGGAAGGATAGG | TAACATGAAATTCCTCATCTTTGCATGGTA | GACAAAGGACTTGGAAGAACGAGAAGAAGGAGACGCACCAAACATTGGCCATAAACAACAAGCCATCCAACAATTACAACTCGCACAAACGGAAGGATAG[A/G]GAGACATATCCGCCACTACCATGCAAAGATGAGGAATTTCATGTTATCCTTGACACGATGATTGCTGATGGTGCAATCAGGCCTGTTAGACCCCACAAAA |

| | | | | |
|---|---|---|---|---|
| SNP_IGA_67137 | GAAGGTGACCAAGTTCATGCTACACAAAGTAAAAAACTTACCGATCCG | GAAGGTCGGAGTCAACGGATTATAACACAAAGTAAAAAACTTACCGATCCT | GGATCTTATCCATGAAAATAGGAGGGAA | AAAATAATTTCTTCTTTTCTTTTCTGTTTTTTTTTTTCAGAAATATAAAGTTCCCTGTTGTTCTGTTACAAATAACACAAAGTAAAAAACTTACCGATCC[G/T]GGGATGTAGAGTCGGCGCTTCCCTCCTATTTTCATGGATAAGATCCCTTCATCAAGTCCCTTGATCACCTGTTCTACTCATTCAGATTAACAATTATTAA |
| SNP_IGA_79455 | GAAGGTGACCAAGTTCATGCTAAGCGCGTTACCATACCCGAC | GAAGGTCGGAGTCAACGGATTCAAGCGCGTTACCATACCCGAT | GGCGGACCTCGGGACGCTT | CTGGAGCCGAACACTCTAGAGACCAGGCCCAGCTCCTTAGCCTCCGAACTCGAAAAGGTCCGACCCGTAAGCGCTAATTCAAGCGCGTTACCATACCCGA[C/T]GATGCTGGGTAGCCTCTGAAGCGTCCCGAGGTCCGCCGTGATCGCCAAGTCCACCTCTTTCACCGAAAAGAACGCGTCCTTGCTGCAATACCTTATGTCA |
| SNP_IGA_86968 | GAAGGTGACCAAGTTCATGCTCATAATGCTGAAGTGCAAACAGGC | GAAGGTCGGAGTCAACGGATTCCATAATGCTGAAGTGCAAACAGGA | CAGATGTAGAAGTTCCAATTGCATGCAT | TAAAATAACATGGCTACTTTGCATTTATACTTCCTCCTCAAAGTTTGATAAATTTTTTTTGCAGATGTAGAAGTTCCAATTGCATGCATCATTCTGATAG[G/T]CCTGTTTGCACTTCAGCATTATGGAACTCATAGGGTTGGGTTCTTGTTTGCTCCTATAGTTCTGACATGGCTTCTGTGCATTAGTGCAATTGGTCTTTAT |
| SNP_IGA_103771 | GAAGGTGACCAAGTTCATGCTTTCTTGAATAGCTGCATCTCTTTCA | GAAGGTCGGAGTCAACGGATTTCTTGAATAGCTGCATCTCTTTCG | GCAACATCAGCCATCAATGAAACAGATAA | TTGCGGCATCTCGCTGCAGGAATGCCATGTCTCGCTCTTGAAGGGCAGCTTTTTTCTCAGAAAATGCCAAATTTCTTTCTTGAATAGCTGCATCTCTTTC[A/G]CCCATTATGGCCATTATCTGTTTCATTGATGGCTGATGTTGCATCAACCACTAAAACAAACAGAGAGATACAAAATTTCATCAGATATGCCTCTCAAGTG |
| SNP_IGA_120784 | GAAGGTGACCAAGTTCATGCTACCCTCATCACTATGGTTGTTATCG | GAAGGTCGGAGTCAACGGATTACCCTCATCACTATGGTTGTTATCA | GACAATGATGAAGAAGAGAATGAGGTTGAA | CCAAAATACCAGTTGATAGTGATTCTGAAGATTCTGAAGATACAGCTGATGACAATGATGAAGAAGAGAATGAGGTTGAAGACGAGGACAATGGTAATAG[C/T]GATAACAACCATAGTGATGAGGGTAAGTGTTAAGTTCATAAATATGTCATGCGCTACCTATAATAAATGACAGCTAGGGCTATGAGGATGAGAATTTTCT |
| SNP_IGA_125219 | GAAGGTGACCAAGTTCATGCTAACTTGCACATTTGAGGCATGAAGC | GAAGGTCGGAGTCAACGGATTCAACTTGCACATTTGAGGCATGAAGT | GGTCATGAAGGTGGAATTTTTACTGTTGAA | AAGACAAAGAACAAAACACCAACAGTCGTTGAAACATAATGTTAACAATGCATGAATACCCTGTAACTGGATCAACAACTTGCACATTTGAGGCATGAAG[C/T]GGGGCTTCAACAGTAAAAATTCCACCTTCATGACCTTGGCCTTGCTTGATATGTTTCTTTACCTGCAAAGACAGAGTACTTGTTCAACATTGGAATAGCC |
| SNP_IGA_123719 | GAAGGTGACCAAGTTCATGCTAAGCAACTGTGGATTGGATGGACT | GAAGGTCGGAGTCAACGGATTGCAACTGTGGATTGGATGGACC | TTTCAGGTTTTTGTGGCTGCAACACTAAA | GTTTGTCGAATATGAGATTGTGTAGATGCATGTAACAAATCTCTTTTGTTATTATCCTTTTCAGGTTTTTGTGGCTGCAACACTAAAGAGAAAGCAGGTT[A/G]GTCCATCCAATCCACAGTTGCTTCATGGAAGCCAAAACAATGTGCCAAATGACACAACCACTTTCTCTACTGACGCTGGCAGTTTTCTTCACACTGAAAA |

| | | | | |
|---|---|---|---|---|
| SNP_IGA_238135 | GAAGGTGACCAAGTTCATGCTCCGATTCAACATGACTAATACGGG | GAAGGTCGGAGTCAACGGATTCCCGATTCAACATGACTAATACGGA | AAAGGCATGTTCCGCAATTTGATCACTT | TTCATCGTCTCATCAATCACCTGCCGCGCATGGTAGTAAGAGCAATTCAAAAGGCATGTTCCGCAATTTGATCACTTGTGGTGCTGTCGACACAAATGAC[C/T]CCGTATTAGTCATGTTGAATCGGGCCAATAAAACAACTTCTTCCGACAAGTCTACCAGATGGAGCAGCCAAGACATTATCAAGGCTGATCCTATTTGTAA |
| SNP_IGA_136625 | GAAGGTGACCAAGTTCATGCTGAAATTGTCAAATGTGGATGTAATGCAA | GAAGGTCGGAGTCAACGGATTGAAATTGTCAAATGTGGATGTAATGCAG | CGGATTTTCATTACTATACTGCAAAAGTTT | GTGCGAAAATGATATACGACGCTGATGATCGTGGCGACGTGATTGATGGTGATCTTGGTGAGCACTTTGATCTGAAATTGTCAAATGTGGATGTAATGCA[A/G]GAGAAACTTTTGCAGTATAGTAATGAAAATCCGAATAGAACAGTTGTGAATCCATACATTCATTTCGGGCAACGGTCAATTTGGCCTAGGGGATTGCCTT |
| SNP_IGA_148760 | GAAGGTGACCAAGTTCATGCTGAGAACCAAATTATCTGCCTGTGGTT | GAAGGTCGGAGTCAACGGATTAGAACCAAATTATCTGCCTGTGGTC | CATCTGATGCCTGAGGTCAACCTTT | CGGCCTGTTGGAGTGCAAAGTCAAGCATCCATTCCTCTGCATTTTTCTTTTCGTCCATCATCTGATGCCTGAGGTCAACCTTTTCTGCTTCAGGATCAGG[A/G]ACCACAGGCAGATAATTTGGTTCTCTTGGATTGTACTTCCTTTGTTCCTCCTCATCCACGCTAAGTCTCTTGAATTTGCTTCCTCTTCGGTAGTTGCAGG |
| SNP_IGA_195134 | GAAGGTGACCAAGTTCATGCTCCAGATGAGATACCCTTGAATGCAA | GAAGGTCGGAGTCAACGGATTCAGATGAGATACCCTTGAATGCAG | CCGGATAGAACTGCATTACAGTTTATGAA | TTGGTTTCTTAATATTATTATCTTTTCATCTCAGGGAACAAGAAAAATTAAAGGCATCATGGTGAAGTTTCCCAAGCCAGATGAGATACCCTTGAATGCA[A/G]AAAGCTTCTTCGGGATGGTAAATCTTGAAATTTTCATAAACTGTAATGCAGTTCTATCCGGATATGTTGAATATCTACCCAACGAGTTGAGGTTCATTGA |
| SNP_IGA_218801 | GAAGGTGACCAAGTTCATGCTGGTCTTCAGTTTCCAGGTTGCG | GAAGGTCGGAGTCAACGGATTTGGTCTTCAGTTTCCAGGTTGCA | AACTCCACTGAAAGATTTTGGATGTGCAT | AGAGGAAGCCTTACCAACTACTGATGGAAACCTTTTGACAGCGGGCATACGAAGGCAAACTCCACTGAAAGATTTTGGATGTGCATCAGTGGGTTTCAAG[C/T]GCAACCTGGAAACTGAAGACCAAAGGAAAAAAAAAAAAACAATTATCATATGAATATGAGCTGAATGAGATGCAACATTGGCAAATTACGAAACAGGAGC |
| SNP_IGA_242118 | GAAGGTGACCAAGTTCATGCTATAGCCAATGCGATGAACACCAGT | GAAGGTCGGAGTCAACGGATTAGCCAATGCGATGAACACCAGC | CTAACAACATCTTGGCCGTCGGATT | TCAGATGGGGTGATCTGCTACTGGACCCAGATCCTAACAACATCTTGGCCGTCGGATTGACGGGGCTGCTGACGTGGGCAAGCGTGCAGGTGCTATGGCA[A/G]CTGGTGTTCATCGCATTGGCTATACTTGTTGCTGCTGTTAAGTATTCTTTCATAGCTGCTGTTCTTCTTTTCATTCTCATTGCCCTTCTTTAAAAATTCT |
| SNP_IGA_252957 | GAAGGTGACCAAGTTCATGCTCCAATAATCCTCCACGTAAGCAACA | GAAGGTCGGAGTCAACGGATTCAATAATCCTCCACGTAAGCAACG | CCTATAACGACGTGCTTTCTTAGGATAAA | AATTCTTCATTACGTAATGTAATTGAGCGATGTTTTGGTGTGTTGAAAGCTCATTTTTCAATTTTGAAATTGATGCCCAATAATCCTCCACGTAAGCAAC[A/G]AAGTATCCCTTGCAAGTTGTGTGTTACATAATTTTATCCTAAGAAAGCACGTCGTTATAGGTTGTTTGAAGAGTTTCAAGTAGAAGACATGATTGTTGAG |

| | | | | |
|---|---|---|---|---|
| SNP_IGA_258854 | GAAGGTGACCAAGTTCATGCTTACACAAGGGGTCTTTGAGCAGTT | GAAGGTCGGAGTCAACGGATTCACAAGGGGTCTTTGAGCAGTC | TCCTATAGGTACTTCCTGTGATGATTGAT | TATGGAGAATTATTTGGAAAATTCCATATTGTTCTTAACCCGTTATGCTCCATCCTATAGGTACTTCCTGTGATGATTGATGTTGGAACTAACAATGAAA[A/G]ACTGCTCAAAGACCCCTTGTGTAAGTGTTTCCTCATCTAATCTTCAATACGCGGTACAAAAAACAGCTATTTTGATTATTTACAATGTTAATATGATGCT |
| SNP_IGA_266063 | GAAGGTGACCAAGTTCATGCTATCATTTGGAGAGTAACCAAATATGATTG | GAAGGTCGGAGTCAACGGATTCATCATTTGGAGAGTAACCAAATATGATTT | AAATGCCAATTCGATTTCACTCAACACTTT | TGTAGTTGGGTTATGGCTCAGAAAATCCTCACATAGTTTTGGGTGAGGAGTTTGGCTACACCATGTTTTCACATCATTTGGAGAGTAACCAAATATGATT[G/T]GAGAGAGATAAAGCACCAAGAAAAGTGTTGAGTGAAATCGAATTGGCATTTTTGGATGGCTTTTGATCAAGAAATGTGTATGAAATATGGGCGATTTGGG |
| SNP_IGA_275502 | GAAGGTGACCAAGTTCATGCTGAGAGCCAGGGAAATTGCCAAG | GAAGGTCGGAGTCAACGGATTGGAGAGCCAGGGAAATTGCCAAA | GCAGCTTAAGAGAAACATCTCAATACCTT | CCACATGCCATACCTCCATTACTAATAAACAGAAGCCAAGAATGACTGAAACATATGCAGCTTAAGAGAAACATCTCAATACCTTGCGTTTAGCAATATC[C/T]TTGGCAATTTCCCTGGCTCTCCACTCATCAGCTTCTCGATCAGCTTCTTCGAATCTTTCGCGCTCCTGTGCATCACCAAAGCAATAAAAAATGTAAGCAA |
| SNP_IGA_283325 | GAAGGTGACCAAGTTCATGCTACAGATTATATGAAGCAGCTTCTTTCG | GAAGGTCGGAGTCAACGGATTCACAGATTATATGAAGCAGCTTCTTTCA | GTGATGCAAGTGAAACCAACCAAATTCTA | TTAAATACACATAATCAGAAATCTTACCCAAGTGGTCTTCATTCAAGTGATGCAAGTGAAACCAACCAAATTCTACTTACATGTGATGAGGGTATTTGAG[C/T]GAAAGAAGCTGCTTCATATAATCTGTGATGTGGTACCCACCGATGTTAGTTCGGCAGCTTCCTTGATACATAGGTTCGCCGTCAATAAACTGGGAATGAA |
| SNP_IGA_285058 | GAAGGTGACCAAGTTCATGCTCACCACTACATTGAAGGAATAGTATGTT | GAAGGTCGGAGTCAACGGATTACCACTACATTGAAGGAATAGTATGTC | GGCTTGGCTTGTCAGCTGAGCT | TGATTGCTCCCATACGCCAATTCAACATCAATGTGCAAACAAATTCCCCATGTACAGTGGGAATACTTGTTGACCGAGGCTTGGCTTGTCAGCTGAGCTC[A/G]ACATACTATTCCTTCAATGTAGTGGTGATTTTCATTGGTGGAGCAGACGACCGTGAGGCGTTGGCCTACGCAGCACGGATGTCTGGCAATCCGGACGTGG |
| SNP_IGA_898654 | GAAGGTGACCAAGTTCATGCTGGTCAGACAATTGAAGAGGCACA | GAAGGTCGGAGTCAACGGATTGGTCAGACAATTGAAGAGGCACG | TCTTTCACTTGTAACAGAGCTTGTGCAT | AGTTGGGCACTAATGGCGGCGAAAGAAGCAATTTTTGTTGAAGCTGCAAATGGATTTGACTTGCAACTGGTAGCACCTGGTCAGACAATTGAAGAGGCAC[A/G]GAGTGGAATCAAAGGGCATGCACAAGCTCTGTTACAAGTGAAAGAGCTGATAGATTTAGAGTCATGGAGAGAAGTACAAATAGCTCTCAGGAAGAGCTCA |
| SNP_IGA_299796 | GAAGGTGACCAAGTTCATGCTGCCACTGGAGAAAGCCTAACCA | GAAGGTCGGAGTCAACGGATTCCACTGGAGAAAGCCTAACCC | CACATATTGCTGCTAAAGTAGCTGCTAAA | GTAGCCAAACCTACAAAAGCCAATTGAAGAATGTTAACTTTTAGGCCTATTCAGTCACAAGAAATCTTACAAGAGCAGAGCCACTGGAGAAAGCCTAACC[A/C]AGTTCATATGATTTAGCAGCTACTTTAGCAGCAATATGTGCCGAAATCTTTCTAATGTTTGTAAATGGAGGGTATATGAGTCCCTTGTCATAGTCTTCCT |

| | | | | |
|---|---|---|---|---|
| SNP_Pp03-3917979 | GAAGGTGACCAAGTTCATGCTGAATCTAGCCAAGAGGCTTGCTG | GAAGGTCGGAGTCAACGGATTATGAATCTAGCCAAGAGGCTTGCTA | GGAGCTTGCTCTATCTGACTGCAA | TCAAAAGTGGATGGAAGTGAACTAGCAGATGAGACATTGTATAGGCAAATGGTGGGGAGCTTGCTCTATCTGACTGCAACCAGACCAGATATCATGTTTG[C/T]AGCAAGCCTCTTGGCTAGATTCATGCATAATCCAACCAAGAAGCACATGGGAACAGCAAAAAGAGTGCTGAGATATGTTCAAGGCACCATAAACTATGGA |
| SNP_IGA_322698 | GAAGGTGACCAAGTTCATGCTAGACGGTAATGGTTGCATGAGCAA | GAAGGTCGGAGTCAACGGATTGACGGTAATGGTTGCATGAGCAG | TCATAGGACTCAATAAGTCCTATAGCGTA | GGCTCGGTAGATGCATGAGACGTTCCAAATTCTCGTTGCTTCTCTTCTTGAGAAATCAAAGAATATGCCTTGCGAACAGACGGTAATGGTTGCATGAGCA[A/G]AATTTGTCCTTGCACTACGCTATAGGACTTATTGAGTCCTATGAGGAATTCCATAAGTGCATTTCTCTCCTCTTGTTCGCTATGTTTCTTCATAGCACCA |
| SNP_IGA_428056 | GAAGGTGACCAAGTTCATGCTGTACGCATCCTATGTTTGGACCC | GAAGGTCGGAGTCAACGGATTGTACGCATCCTATGTTTGGACCT | GACCTTGTCATAAACAAAAGGAAGACCAT | TTGTTGATGTCCTTTCGGTAAAAGAATTTCCAAGAAATCTGTTACTTCAAAATTTGCCGCTGGATTGTGATATACTCTGTACGCATCCTATGTTTGGACC[C/T]GAGAGTGGAAAAAATGGGTGGAATGGTCTTCCTTTTGTTTATGACAAGGTCAGGGTCGGAGGTGACGAATCAAGAGTGTCACGGTGCAACAAATTTCTGG |
| SNP_IGA_894039 | GAAGGTGACCAAGTTCATGCTCCTCTTATTGTGGTGCTCCC | GAAGGTCGGAGTCAACGGATTAACTCCTCTTATTGTGGTGCTCCT | CTAAACATTTGCGAACAGATAAAGACGCTA | TGAACTCATAAGAGAATCATTACATTCCTTCCCAGACTTGGGCCATCCGGGCCCATGGACTAGTGGTCCCGAAACTGAACTCCTCTTATTGTGGTGCTCC[C/T]ATAAATCTTTTCTAGCGTCTTTATCTGTTCGCAAATGTTTAGTTGGATCAACCAGAAAACCAGAAACAGCTTCCTCCCTGTGAGGGTTGAACATTTCACT |
| SNP_IGA_325850 | GAAGGTGACCAAGTTCATGCTTACGCATTGTGCGGAAATAATGCAAA | GAAGGTCGGAGTCAACGGATTACGCATTGTGCGGAAATAATGCAAC | CAATACTCCGAGCCTGTGGACATAT | GAACTTGACCTGACCTCCCTATGATATCCACCATACATGAGTAATGCTCCAATTGAGGGTGTAAACCATAGTCATTACGCATTGTGCGGAAATAATGCAA[A/C]CCCTTTTCAACTTGTCCTATATGTCCACAGGCTCGGAGTATTGAAACAAAAGTGGAGTGGTTTGGCTTAACATTCTCAAGCTGCATGTTCTCAAAAATTC |
| SNP_IGA_344612 | GAAGGTGACCAAGTTCATGCTTCCATCATGAAATATCACTGGCTTT | GAAGGTCGGAGTCAACGGATTCCATCATGAAATATCACTGGCTTC | ACAGCAAGAGTGGTTCAAATTCAGTGAT | AATTAATTATGTGTATTGGATTGTAACAGGAAGCCTATGTAGTGAACAAGGAAGACAGCAAGAGTGGTTCAAATTCAGTGATGCCAAGGCAAAAATACCC[A/G]AAGCCAGTGATATTTCATGATGGAAGGCTAGCATTCTTGCCTACACCATCTGCAACTCTTGCCATGTTCATGTGGCTCCCTCTTGGAATAATCCTAGCCA |
| SNP_IGA_349233 | GAAGGTGACCAAGTTCATGCTCCGCCATCTTCGCTGTCCG | GAAGGTCGGAGTCAACGGATTATACCGCCATCTTCGCTGTCCA | CGTTTCTTTCCATTCCCGAGCGAAT | GAGGCCCAGGCGCTCGCCAGCAAGCGTCGGCGTATCAGTATCGCACCCTCTCCAAACTCCGCCGTTTCTTTCCATTCCCGAGCGAATCACAAAGAAGGCT[C/T]GGACAGCGAAGATGGCGGTATTTACACCGCGGCTCTCAAAGGAAGCCAAAGAGCAGCCCCCGAGCCTCCACGCGGTGGATTTTCGACGGGGAGAGTCAAT |

| | | | | |
|---|---|---|---|---|
| SNP_IGA_356179 | GAAGGTGACCAAGTTCATGCTCAGTAGTCAGGATAAATGCATGTGTT | GAAGGTCGGAGTCAACGGATTCAGTAGTCAGGATAAATGCATGTGTC | TTTCATGCGAGGCTCAGAAACAAATGAA | CACTGCCTGCAGAATATGACCGAGCAGACCTTGACGTATACCTCTCCATTCCCCGAACTTTCATGCGAGGCTCAGAAACAAATGAACCATGGTCTTTAAC[A/G]ACACATGCATTTATCCTGACTACTGGAAGCATAGTTCCTTTTACTCCTTTTATGACAGTAGTCTTAATTTGAGTCGTTCTGCTCACACCACTACAATCAC |
| SNP_IGA_368926 | GAAGGTGACCAAGTTCATGCTAGCCCGTTATTCACACCTCTATTC | GAAGGTCGGAGTCAACGGATTGAGCCCGTTATTCACACCTCTATTT | GTTGGCATTGCAAAGAGGGCATGAT | TATAGCTCTGATTCTCCTCATCTCTTTCTTTAAATAAGTTGCTGACCAGTATATACCAAGTCAAATGATTTGGAACGAGCCCGTTATTCACACCTCTATT[C/T]AAAAGTAAACAAGCATCATAAATCATGCCCTCTTTGCAATGCCAACTGATCAAAGTGTTATAAGTAATAACATCAGGCCACATCCCTTCAACTTGTAACC |
| SNP_IGA_379856 | GAAGGTGACCAAGTTCATGCTGTTTGCTTCGATCTGGTCATCC | GAAGGTCGGAGTCAACGGATTGTGTTTGCTTCGATCTGGTCATCT | AATGAACCCAAAAATATCAGATTTCGGCAT | ATTTTGTAAATCAGTTGTGCATGTTTCGGATTGAGGGCAGAGAGTGAGAGTTTTACTATGTGCCAACAACTCTGTTTGTGTTTGCTTCGATCTGGTCATC[C/T]CCGAACATTCTTGCCATGCCGAAATCTGATATTTTTGGGTTCATTGAGGCATCCAGTAAAACATTGCTTGCTTTCAAATCCCTATGGATAATTTTTAGTC |
| SNP_IGA_389204 | GAAGGTGACCAAGTTCATGCTAGAGTCTGAAATTCCCTTCCCTCAT | GAAGGTCGGAGTCAACGGATTAGTCTGAAATTCCCTTCCCTCAC | CATGGGGATCTTGCCATGTAGTCAA | AATTGTAAAGCTTCCTAATCCAATCCATGTGCTTTGCCTCATCTTCATGGGGATCTTGCCATGTAGTCAAGTACTGAATTTTAAAGAGAATTCCATTTCT[A/G]TGAGGGAAGGGAATTTCAGACTCTGAAATTCTGCTCATCTGTCCTCCATATGGATTGAAAATCACCAGAGGGCTCTCTTCCTCCATCAGCCTTTTCCAAA |
| SNP_IGA_395253 | GAAGGTGACCAAGTTCATGCTCAGACATCATCCTCTTCACG | GAAGGTCGGAGTCAACGGATTGCTCAGACATCATCCTCTTCACA | GTAGGCCAAGGATCAAGCACATTCAA | CGTTAGGATGTACAGAATTAACGAAATAAATCAATGTTAACTTGTGGCTGCTTCATTGTAGGCCAAGGATCAAGCACATTCAAAACCTAGTTCAAGAAAA[C/T]GTGAAGAGGATGATGTCTGAGCAATTACAGATTCCTTCCACTGCCTCTGGTCGTCAATTAAAACCAAATTCTGATAATCGCTTCACCAAACCAAGCTTTA |
| SNP_IGA_398213 | GAAGGTGACCAAGTTCATGCTTCCTTTCCAAAATCTTAGCTCTTGTA | GAAGGTCGGAGTCAACGGATTCCTTTCCAAAATCTTAGCTCTTGTG | ATGGGTTGAACTCAAGTTCTGAGAAAGTT | AATCTCCATGATGATGAAGATCAGCAGAAGCACCATCAACACCTACACTAGCACGCTTAAAGCTCTTGGACTTCTTCCTTTCCAAAATCTTAGCTCTTGT[A/G]GTACCTCCTGGGGTCACAACTTTCTCAGAACTTGAGTTCAACCCATTAGGATCATTTCCTCTCTTTATTGCTGGTGGCCTTGGAGACTTTGACTTCGGAG |
| SNP_IGA_403741 | GAAGGTGACCAAGTTCATGCTAAAGACCTGGCTTTTGAGTTGCCT | GAAGGTCGGAGTCAACGGATTAGACCTGGCTTTTGAGTTGCCC | ACCTCAACTTCAAAACCACCAGTGTTTTA | TCTTGAGACTTGGGAAACACCCTGACGCAGGAGGAGCATTGGTAATACAGTTTGACCTCAACTTCAAAACCACCAGTGTTTTACACATGAAAATGAATTG[A/G]GGCAACTCAAAAGCCAGGTCTTTATCTGTTGTTTCAAAACAAAGATCAAGTTCAACAACATTATGCTGGATGGCAGTTCGAATCCAACCATCAATACCAG |

| | | | | |
|---|---|---|---|---|
| SNP_IGA_409453 | GAAGGTGACCAAGTTCATGCTCCGATATCTACAGCTTTGGAGTC | GAAGGTCGGAGTCAACGGATTGTCCGATATCTACAGCTTTGGAGTT | ATGGGTTTCCTCCCGCAAGCAATTT | CACAAACAACAAATGTTGCTGGAACCAGGGGTTATATGGCTCTGGAATACGTTACCACAGGAAAGGCTAGCATGGAGTCCGATATCTACAGCTTTGGAGT[C/T]GTTGCTTTGGAAATTGCTTGCGGGAGGAAACCCATTGATCTCAGTTTAGAAAATAGCCAAATCGAAATGGTGGAGTGGGTGTGGGAGCTTTATGGAGAAG |
| SNP_IGA_420819 | GAAGGTGACCAAGTTCATGCTGTAGACGACACCATTGACGACCT | GAAGGTCGGAGTCAACGGATTAGACGACACCATTGACGACCG | CGACGTATATGGGTTTGGATTTGATTCTT | AAGAAACTACCAGAAATACCAACAGAATATCTAATCGATGACCTCTACTACTTTTGCGACGTATATGGGTTTGGATTTGATTCTTCAACTCATCAACACA[A/C]GGTCGTCAATGGTGTCGTCTACAGAACAAGTATCCGTGATGATGATGAAGGAGGGGTTCAGTTTAATGTGTATACATTGGAAACTAATTCTTGGCGACAG |
| SNP_Pp04-13754607 | GAAGGTGACCAAGTTCATGCTCAACGTGTTGTATTTGATTGAATACCATT | GAAGGTCGGAGTCAACGGATTCAACGTGTTGTATTTGATTGAATACCATC | GACTATAAGGTTTAGCGCTCTCGTCTT | CGAATTTCATGCCAAAAGTTTATATGTATTTTTTTAAGCTTATGAATTTAAGACTATAAGGTTTAGCGCTCTCGTCTTTTCATATAAATATTATTGTTTA[A/G]ATGGTATTCAATCAAATACAACACGTTGCTATAAGGATTATGATTTTGTTTTTTCATAAAAATTAAAATTGCTTTGGTTGGAGTATGACCAAACTAGAAG |
| SNP_IGA_440116 | GAAGGTGACCAAGTTCATGCTGAACATTTCTATACTGCCACGACC | GAAGGTCGGAGTCAACGGATTCGAACATTTCTATACTGCCACGACA | CTGTGCAATGAAGTGCAGCAACACTT | AATGAAGGAAAGAACATGTGGTGCATGGTCCAGGACTTTGTCCTTGGGGGCATGCGTGTACTGTGCAATGAAGTGCAGCAACACTTTTGAGCGCACCATC[G/T]GTCGTGGCAGTATAGAAATGTTCGGCCTTAACACTCTTATCTCATCATCTCATGCTCAATAAGATCATCATTCTCTCTTTATTAGTGGATATCCAATCGT |
| SNP_IGA_454568 | GAAGGTGACCAAGTTCATGCTGTCCTCTTCTTCTCCTTCTCCTTT | GAAGGTCGGAGTCAACGGATTCCTCTTCTTCTCCTTCTCCTTC | GAGTGGAAGAAGAGGAAAGAACATTTCTA | TCCATGTCCAGTCCAGTGAGCGACACAAAAAGCGGCGCCTAAGAGCAGAGTCAGAGTGGAAGAAGAGGAAAGAACATTTCTATAGCAGATGCGGCAAATG[A/G]AAGGAGAAGGAGAAGAAGAGGACATAGTTTGCTTAGATGAATCTTTCTTCATCGATGACAAGTAAGAGGAAGATACAGTCTTTTTAAGTTTTGTGGTTAT |
| SNP_IGA_480804 | GAAGGTGACCAAGTTCATGCTACCATTTACATCACCTATTGTGTTCGA | GAAGGTCGGAGTCAACGGATTCCATTTACATCACCTATTGTGTTCGG | TCAATCAGACAACTGGGTTCCGGTA | TAGATGCATATGTTGTGAATGTTCCTAGCATGATGAGCATAGATTTCAAGAATCCCTGAAAAGTGATCAGAATCACCATTTACATCACCTATTGTGTTCG[A/G]ATTCAGGTTGTACCGGAACCCAGTTGTCTGATTGAATGATTCCATCAGATCCGAATATAGCTTCTTCCCCTTGAGGATATTCTAATATTTTCTGTTTGGT |
| SNP_IGA_513502 | GAAGGTGACCAAGTTCATGCTAGTATCTTCAAGTTAATAGGCCTGGT | GAAGGTCGGAGTCAACGGATTGTATCTTCAAGTTAATAGGCCTGGC | TTAGTCCATCGTTGGACCTGGTCTT | TGGTGATGGCATTCCATGCACCGTCCACGTTGATTTTGAGATGCCTCATGGGTGGTTTAGTCCATCGTTGGACCTGGTCTTGGTTGGAACAAGGTGTCCC[A/G]CCAGGCCTATTAACTTGAAGATACTCAAGTAGTCTATTATGTGCAATATGGCCTACCAGGCTTGGGCATGAACTTTTATTGTTCCATGGTAAACCATTTC |

| | | | | |
|---|---|---|---|---|
| SNP_IGA_528949 | GAAGGTGACCAAGTTCATGCTGTCACATATCTCCCACTCCCACT | GAAGGTCGGAGTCAACGGATTCACATATCTCCCACTCCCACC | CATCAAAACCTAGGAAACTTTGTTGGGAA | GCGCATAGGTGGTCCAGGATGAGGCATTAAAACAGATTGAATCTAGCCTCTGTGCAATCCATCAAAACCTAGGAAACTTTGTTGGGAAACAAAAGCAGAT[A/G]GTGGGAGTGGGAGATATGTGACAGAACTTTCGCAACCAATCGTATTTTTGGTTGGAATGGCAGTGAGGTCAGCTGAGCCGTAAAGAACATTATTATTACC |
| SNP_IGA_544495 | GAAGGTGACCAAGTTCATGCTTCTCCACCACTTCCTCTACC | GAAGGTCGGAGTCAACGGATTACTTCTCCACCACTTCCTCTACA | GCCGGAGAGAGATACATATGCACAT | GGACGCTCGTGGATGTTCGTGATTTAGCAGAATCACTGCTTCTTGTATACAACAAGTCTGAGGCCGGAGAGAGATACATATGCACATCGCACGAGATCGG[G/T]GTAGAGGAAGTGGTGGAGAAGTACTTGAGGCCTGCATATCCTAACTACAACTATCCCAACAAGTAAGTTTTTCTTACCCTCCAATAATTAGTTTAGTTGG |
| SNP_IGA_548037 | GAAGGTGACCAAGTTCATGCTTTTCATTCGATAGTGCCCTGC | GAAGGTCGGAGTCAACGGATTCCTTTTCATTCGATAGTGCCCTGT | CAAGCCTCTAACACTTCCAACCTGAT | TGGCAGCCATCGGAATACGAGTATTACAGTGGATAACGTATCATGGCTTAGCGTTGAATGTCACCACTGACTTAACCCCTTTTCATTCGATAGTGCCCTG[C/T]GGGTTACGAGACTATCAGGTTGGAAGTGTTAGAGGCTTGCTGAAGGAATTTCAGTCATCCACTGACTGCGAAAGAGCACGTCTACCTGATCCTGATGATG |
| SNP_IGA_553456 | GAAGGTGACCAAGTTCATGCTAAGTTCATATTCTACTTGGCTGAGGA | GAAGGTCGGAGTCAACGGATTAGTTCATATTCTACTTGGCTGAGGG | GAGTTTGATTATGTTTGCGAATTGCCGTT | ATGCAACTATTTCAATAAGTTACCTAGAGAAGTAATATTGGCTTCCCCAATATCAAGCTCTTCGGTTTTCATCCTAAGTTCATATTCTACTTGGCTGAGG[A/G]TTGCTTTCCAGAACGGCAATTCGCAAACATAATCAAACTCTATCTTCTTCTCGATTTCAGATTCTAACTGTTTGAATTTGTTGTCATATGACTGCAACTG |
| SNP_IGA_571548 | GAAGGTGACCAAGTTCATGCTAAGGTCAATTCCAGCACTCTTGGTT | GAAGGTCGGAGTCAACGGATTGGTCAATTCCAGCACTCTTGGTC | TGCAAATTGCTCACCAGAAGGGCTA | GGGTTGTATTAAACTTTACCTGCATGGAAATGAAAGACAGAGAACAGCCTGCACATGCAAATTGCTCACCAGAAGGGCTAGTGCGGCAGGTAAAAATGGC[A/G]ACCAAGAGTGCTGGAATTGACCTTGCAGGAGAGAATGCATTGGAGAGGTATGATACTGGTGCATTTGAACAAGTTTTGGCAACAAGTAGATCAGATTCCG |
| SNP_IGA_572589 | GAAGGTGACCAAGTTCATGCTGTTTACCACTGGGATCTGAGAACA | GAAGGTCGGAGTCAACGGATTTACCACTGGGATCTGAGAACG | AAAGCGGTACCATTTATGCAACCTTCAT | AGATGAATGGAACTGTTCGCTCTTTATCTTTTGCCAATGATGGGAAGCAATTATTGAGCTCTGGTGGTGATGGACAGGTTTACCACTGGGATCTGAGAAC[A/G]GGGGCCTGCTTCCACAAGGCACTTGATGAAGGTTGCATAAATGGTACCGCTTTGTGTACATCTCCAAATGGGACCATGTTTGCTGCTGGTTCAGACAGTG |
| SNP_IGA_585810 | GAAGGTGACCAAGTTCATGCTCAGTGCAGCTGCTTCAGATTCTG | GAAGGTCGGAGTCAACGGATTCAGTGCAGCTGCTTCAGATTCTA | GCAGAAGCATGCAGATACGAAGAATTTAT | GTTTTATCCGGAAAACTACAACTCAGTAGTTCAATGCGATTGGCTGCAGAAGCATGCAGATACGAAGAATTTATCGATACCAACCTGAAAGGGAAATTCT[C/T]AGAATCTGAAGCAGCTGCACTGGCAAAAATCGCGCTTGTATGCACCCACGAGCTTCCCGACCACAGGCCAACAATGCAGGAAGTAATTCTGGAGCTGAGT |

| | | | | |
|---|---|---|---|---|
| SNP_IGA_591439 | GAAGGTGACCAAGTTCATGCTGATGTGAGCATGATTGGGCAA | GAAGGTCGGAGTCAACGGATTCTGATGTGAGCATGATTGGGCAG | CACAATCTCCCTCTGCAGAGTAGAA | TGTTAGATTTAGTAAACAACTTGGGGACAATTGCAAGGTCTGGGACAAAGGAGTTTATGGAGGCATTGCAGGCAGGGGCTGATGTGAGCATGATTGGGCA[A/G]TTTGGTGTGGGTTTCTACTCTGCAGAGGGAGATTGTGACCACAAAGCACAATGATGATGAGCAGTATGTGTGGGAGTCTCAGGCTGGTGGTTCTTTCACC |
| SNP_IGA_595126 | GAAGGTGACCAAGTTCATGCTGAGACCCAAAAACTGCCGGTAAG | GAAGGTCGGAGTCAACGGATTAGAGACCCAAAAACTGCCGGTAAA | GTTCGGAGACTTGAAAATTCCGCGAT | ATATCTTCTCGAACCGGTCAAGTGGGTCCTCCTTGTTCTTCACGCCGAAAACGCCGTCGTTCGGAGACTTGAAAATTCCGCGATTGGGGTTGGAGTTTTC[C/T]TTACCGGCAGTTTTTGGGTCTCTAAAGTTGAAGGCTTTGGGCGGGGGGAGCGGCGAGAATCGGAAGCTATCGGCGTCGAGGCCGGCCATGAGCTCCCATG |
| SNP_IGA_596393 | GAAGGTGACCAAGTTCATGCTGGCATAGAGTTCAGCAGAGAGG | GAAGGTCGGAGTCAACGGATTTGGCATAGAGTTCAGCAGAGAGA | GCACTCAACCCTTTAACTGAGGATCTT | CACCGAGAGAGCACATAGAAGAGATAAGGAAGAAAAAGTTCTCCATAGGAGCAGATGCACTCAACCCTTTAACTGAGGATCTTCACCAGGCTATCAAGAC[C/T]CTCTCTGCTGAACTCTATGCCAAAGATGTTCACTTCCTCATGGAACTCATCCAGGTTGTTTATTTTAAATATACATTCCTACATATCATGACTTAAGTTT |
| SNP_IGA_600493 | GAAGGTGACCAAGTTCATGCTAACCAATTGATGCAAGAAAATAGTGAGAAT | GAAGGTCGGAGTCAACGGATTCCAATTGATGCAAGAAAATAGTGAGAAG | CATCCATATTATCCACTTCGTCTTCAGAA | CATCCTGTGTGGTATCACTACCTTCTCCTAGAGTGACCTCATCAGGCTGAATTTCATCCATATTATCCACTTCGTCTTCAGAACTGTCATCAGAATCTGA[A/C]TTCTCACTATTTTCTTGCATCAATTGGTTCACAGAATTGTCATCCTCTATCACTTCCTTAGCAACATCAACATCCATGACTTTAACTTTAAGTCCTGGAA |
| SNP_IGA_602605 | GAAGGTGACCAAGTTCATGCTGGTCTCTATCGAGGTTTTGGGG | GAAGGTCGGAGTCAACGGATTGGGTCTCTATCGAGGTTTTGGGA | CAAAGTACATGCCTCGATACAGTGTAAT | GAGATGCTTACCTCCAAAGACCCAACCAAAACAATAGGCTTCATGGTGTCATAAATCCCAAAGTACATGCCTCGATACAGTGTAATCCCAATAATTGAAA[C/T]CCCAAAACCTCGATAGAGACCCGCTATTCCATCACTTGACAAGGTTTTACTATAAACATCTAGAATCCCTCTGAACTGGCGCTGACCATTAACTGAAGAC |
| SNP_IGA_616458 | GAAGGTGACCAAGTTCATGCTAAGTTGGAATGTCAGAGGTTTGGGT | GAAGGTCGGAGTCAACGGATTGTTGGAATGTCAGAGGTTTGGGC | TCTCTCAATAAATCATTGAGAGCTCTGAAT | CTTTCAATTGCACACTAGTACATTTAGTCTCCATAAGGAAAATTACATGGGGATTTTTCTCTCTCAATAAATCATTGAGAGCTCTGAATGTCCGAGGATT[A/G]CCCAAACCTCTGACATTCCAACTTAAGATATTCAATGCCTTTGTGCCCAACATCTTTGTGTCCACCACGGATTCATACATGACAATCTCAGTATTTTTTC |
| SNP_Pp06-2176846 | GAAGGTGACCAAGTTCATGCTCCATTGTAGCAAACCAGAATAACAATTG | GAAGGTCGGAGTCAACGGATTCCATTGTAGCAAACCAGAATAACAATTA | CATTGTATTGCCTGATTATTGTGGTTGTAT | ATATTGATGTTATATTGCTGTTATATTGTCATTGTATTGCCTGATTATTGTGGTTGTATTGTGTTAATGAAATAATTTTTTTCATAGTCAGAAATGGAGA[C/T]AATTGTTATTCTGGTTTGCTACAATGGAAAATGGGTAACCTCAAAGAAGATATGCAAATACGAAGGGGGTGACTCAAAAGGCTTAATAGTTCCACGAACC |

| | | | | |
|---|---|---|---|---|
| SNP_IGA_610889 | GAAGGTGACCAAGTTCATGCTAGCTTGAGGAAGAAAGTTGTATTATGC | GAAGGTCGGAGTCAACGGATTAAGCTTGAGGAAGAAAGTTGTATTATGT | TTCATCATTGTTTTCTGTGAACAAACCGTT | TGATTTATCAGCGCAGGAAAGTGTCAGCAAAGTTGAGGGAAGAAAAGCAGAATCCCAGCGCAGCGAAGAATTTAAGCTTGAGGAAGAAAGTTGTATTATG[C/T]ACAACGGTTTGTTCACAGAAAACAATGATGAAATGGGTGCTTTAACATCGAAGAGCAAGAACAGTGAATTTAAGGAAGCAAAAGATGCAGCTCCAGGACT |
| SNP_IGA_627328 | GAAGGTGACCAAGTTCATGCTGAGGTTTCTAGTAATCTATTTTCTCTTTG | GAAGGTCGGAGTCAACGGATTCTGAGGTTTCTAGTAATCTATTTTCTCTTTT | ACTCATGCTATGTCTTCTTTTGCAACCAA | ACAACTCTTTCATTTATTTTTATTTTTCAGTCTTTCACTACAAAAGAACTCACATGTATTGTGAAGTTCACTGAGGTTTCTAGTAATCTATTTTCTCTTT[G/T]GGGGGCAGTGCATTACACTCGTTGGTTGCAAAAGAAGACATAGCATGAGTATTACAAGTGGACCTTTTTCGCCTTGGATACGGATAAATGACGGCCAACT |
| SNP_Pp06-7216093 | GAAGGTGACCAAGTTCATGCTATATTTCTTATTTAGGGATATAATTGGAAATAA | GAAGGTCGGAGTCAACGGATTATATTTCTTATTTAGGGATATAATTGGAAATAC | TTCCCATCCAAATCCTAATTAATTTGGAAA | TTATGACCAATTTACCACTAGTAACAATTAAAAATACCAAGGCAAACATATATTACACTTTTTAATGCATATTTCTTATTTAGGGATATAATTGGAAATA[A/C]ATTTTGTTTGGTTTTCCAAATTAATTAGGATTTGGATGGGAAATACAATTTACCACTATAGGATTTGGATTGGTGTCGAAAAAAGGGATTTCTATTAACA |
| SNP_IGA_639062 | GAAGGTGACCAAGTTCATGCTGGTCGCGAGTGTTGTTGTCCTT | GAAGGTCGGAGTCAACGGATTGTCGCGAGTGTTGTTGTCCTG | GCACAGCACAAGGCAGAATCATGAA | GAAGTTGCTTGCATTGCAAAAGCAAAAGCAAAGCTAGAGTTTCAGCGAAAGCACAGCACAAGGCAGAATCATGAACCGTCTGAGTTTGGAGCAGGTGCTC[A/C]AGGACAACAACACTCGCGACCCCAACTCCATTTCAAGTCTCAAGCTCAATCACAAGGCTCTTTCGGATGTACGATGTCGTTTTTCTGCTCTCCCCCTATC |
| SNP_IGA_640395 | GAAGGTGACCAAGTTCATGCTAAACTTAAGAAGCGGAGGCCAATG | GAAGGTCGGAGTCAACGGATTGAAACTTAAGAAGCGGAGGCCAATA | TATTCCTCATATCCAGCAGGGCCAT | TGTAGGCAGCTTCCAATATCTTGAGGTTTTGGGTATGTGCTTCTTCTGGGAAGCTGTAATCAATGTATTCCTCATATCCAGCAGGGCCATCTTTGGTTAC[C/T]ATTGGCCTCCGCTTCTTAAGTTTCTTTGGAAGCTTAGACTGGACTAAACTAACATCACCGAGATCTCCGAAGCCAGCCTCCACATTCAACATTTCTTCAA |
| SNP_IGA_655825 | GAAGGTGACCAAGTTCATGCTAGAGGATGTATGTGGTGAAGAGG | GAAGGTCGGAGTCAACGGATTGAGAGGATGTATGTGGTGAAGAGA | TCAAGCGCGCCGTTATCTTGTCAAA | CCACCAGCACAGGGTCGCAGTGGTCGCTGCTAAGCCCATAGCTCAGCTTCTTCAAGCGCGCCGTTATCTTGTCAAAATGCACCGCCTCCTGCCGCCCATC[C/T]CTCTTCACCACATACATCCTCTCTCTCTATCTCTGTCTCTGAACGATTTCGTCGAACAGGTCTTGTGCAAGAGAAAGTGAACAAGGAGGAGACTCTGGGA |
| SNP_IGA_663057 | GAAGGTGACCAAGTTCATGCTAAGGGTCCTACATAATTCTCAAATCCA | GAAGGTCGGAGTCAACGGATTGGGTCCTACATAATTCTCAAATCCC | GAAGACTATCAATGGAGATGACCTTCTTT | AAGGGTCTTCTTCTTGTCTAGTCATGGAGTTCTTCTCCCCCTCAGTCTCCCTATACTTGTTGAGATACCCTTTCAAGGGTCCTACATAATTCTCAAATCC[A/C]AGGGTTGTCATGGCCCAAAGAAGGTCATCTCCATTGATAGTCTTCCTTTTCTCCCTCAGGCACTTGTCAGACGCTTCACCCGTGATGAAGCTTATGAACT |

| | | | | |
|---|---|---|---|---|
| SNP_IGA_671806 | GAAGGTGACCAAGTTCATGCTGGGGTTTTGAAGTACTTTACCTATAAG | GAAGGTCGGAGTCAACGGATTATGGGGTTTTGAAGTACTTTACCTATAAA | GGCCGTGCAACTTGGAGCAGTT | GTCAACAACCTTGACAGAAAATGTACATACAATTAACATAACGAAGGTGGCAAATCGCAGGGCCGTGCAACTTGGAGCAGTTTTCTTAATCTTCTTCTCG[C/T]TTATAGGTAAAGTACTTCAAAACCCCATAAGTTTGAATTGCTGTCAAAAGCACAAAGTAATGTTTTATTGTTGTGTTAAATTATGCTTTGTATTGGATGC |
| SNP_IGA_682254 | GAAGGTGACCAAGTTCATGCTGAAGTGGAATTCAGCAGTCTTTCG | GAAGGTCGGAGTCAACGGATTGAAGTGGAATTCAGCAGTCTTTCA | GACTTGCTCTCTTTAGAAGATGAACTGAA | CAGAACATCCTTCACATTTTGACCAGCAGGCCATCATGAAGCCTAGGATCAGCTTCTCCGACTTGCTCTCTTTAGAAGATGAACTGAACAATGCGGATTT[C/T]GAAAGACTGCTGAATTCCACTTCAAGTGGAGAAAACTCACCGAGCTCGCCATTGATGTGTTCATGGCCAGGCAGCAGTCCAATGCAACAACGTCGTTACT |
| SNP_IGA_691624 | GAAGGTGACCAAGTTCATGCTCCAGAACACCCCAAAACCCAGA | GAAGGTCGGAGTCAACGGATTCAGAACACCCCAAAACCCAGC | TAATGCAGGATTTTTCTGCATATGACGAAT | GACTGTGAGCTGACGCCTCCAAGCATATCATCTTGCAGAAACTCACCATCCTCATCAAGAATGGCTTTAGCTTTGCGAGCCAGAACACCCCAAAACCCAG[A/C]TTTTGTGGCCGATTCGTCATATGCAGAAAAATCCTGCATTATTTATAAGGCATTATCAAAATCATCTCTTACCACACAAATTAAGAACAGGGTCACAAAC |
| SNP_IGA_696341 | GAAGGTGACCAAGTTCATGCTTCCTTCTCCATTTCAAAACACCCG | GAAGGTCGGAGTCAACGGATTTTCCTTCTCCATTTCAAAACACCCA | CTTTAAGTGCTTGATGGGCAATAACAGTA | GGAGATTGCCAGATGATAGAAATGTGAGGTGGAGCAAATATCTGTGCAGGAACTTTAAGTGCTTGATGGGCAATAACAGTAGGGGTTATTCTAAGTGTGT[C/T]GGGTGTTTTGAAATGGAGAAGGAAAAACTTAAATGGGTTACAAACAGCTCCCTTCCCATTGATTTTCTGATCAATGATGTGTTGGCTATTAAGCTGGGGG |
| SNP_IGA_700653 | GAAGGTGACCAAGTTCATGCTATTCAGGAACAGAGGACAAGGCT | GAAGGTCGGAGTCAACGGATTCAGGAACAGAGGACAAGGCG | TATCTCCATCTAATTTACCAGTGGGGAAA | TAATGATGTATGAGTGGACATTCACAATCTTCTTTATATTATTTGGAGGTTATCTCCATCTAATTTACCAGTGGGGAAAGGGAACAGCAGTGCAAGCAGG[A/C]GCCTTGTCCTCTGTTCCTGAATCAGTTGATTATGATTCTGGGAAAGGTTAGTTGGCTTGAGTTAATCCAATGCTGATGTACTAAATTTCATCGAATTAGT |
| SNP_IGA_726222 | GAAGGTGACCAAGTTCATGCTAATGACCCTGGAAATGATATAATGGTAATA | GAAGGTCGGAGTCAACGGATTGACCCTGGAAATGATATAATGGTAATG | GGGACTCTTGCATTTAGTATACATGTCAA | CTGCTGGAGCACTGATTCATGGAAAGGAAACTCATTGTTATGCAATAAAATGGATCCTGAACCTAGACAGAAATGACCCTGGAAATGATATAATGGTAAT[A/G]AACGGTCTGATTGACATGTATACTAAATGCAAGAGTCCCAAAGTTGCACGAATGATGTTTGATTCTGTTGCACCAAAGAAAAGGAATGTGGTGACTTGGA |
| SNP_IGA_717591 | GAAGGTGACCAAGTTCATGCTGGATTTGCATCAAATGGTGTGAGC | GAAGGTCGGAGTCAACGGATTCTGGATTTGCATCAAATGGTGTGAGT | TCCCCAACTTTACTATAGCAGTTGCAAA | AGCTTCTTTTGAAGAGAGGTATCTTGCAGATCGACCAGGAGCTTGCTTCAGATAGATCTACGACTGGTATTGTTTCTGGATTTGCATCAAATGGTGTGAG[C/T]TTCAGTCAGAGCTTTGCAACTGCTATAGTAAAGTTGGGGAGTCTCCAGGTTCTTGTCGGAAATGCCGGAGAAATTAGGAAAAAATTGTAGAGTTTTTAATC |

| | | | | |
|---|---|---|---|---|
| SNP_IGA_704075 | GAAGGTGACCAAGTTCATGCTGGGTGGGAGACAGTGCATAGAT | GAAGGTCGGAGTCAACGGATTGGTGGGAGACAGTGCATAGAG | GGGTCTGTTGGGTTTAGCTTGGATT | GCTTCTAATCCTTTGCTTTTTAATGGGAAAAAGTGGGCGAATGGGTCTGTTGGGTTTAGCTTGGATTCGTCAATTGTGTAATAATGCTTTTTTGGTAGAA[A/C]TCTATGCACTGTCTCCCACCCATCATCAGCCGTGTCTCTCCTCCCACCTAAACCACCACCACTCCACCCAAATCAATACTATAGCTACCACCATTTCAAG |
| SNP_Pp07-5647370 | GAAGGTGACCAAGTTCATGCTAAGAAGATGAAGAAGGCGCGGTA | GAAGGTCGGAGTCAACGGATTAGAAGATGAAGAAGGCGCGGTG | CCGTCTGTGATGAAATCAACCAGCT | TAGCTGGCCCGTCGGCGCTGATATCGAGCTCCACGGCCACGGCCCTGGAGCTTCCAGGCCCATTCACAGAAGAAGATGAAGAAGATGAAGAAGGCGCGGT[A/G]ACGGGGAGCTGGTTGATTTCATCACAGACGGATTGAAGGCGGTCGATGCGGCGAGCCGCGGCGACGATCCTGCATCCAGCTTTAGCCAAGTCCAAGCATA |
| SNP_IGA_752104 | GAAGGTGACCAAGTTCATGCTGCCTTACTTGAGTACCTGAAAGC | GAAGGTCGGAGTCAACGGATTAGGCCTTACTTGAGTACCTGAAAGT | GAAGTACGTAGTCCTTTTAGTACCCGT | CGTTGATGGGTGTTCGACGCTGGAATTCATATACAAATATGAAAAGTTAGCCTCAATTCTGGTACCGCTCCATGTGAGGCCTTACTTGAGTACCTGAAAG[C/T]GCACGGGTACTAAAAGGACTACGTACTTCAGCCTTTCAGGTACTAAAAATACATTTGCTGCTTTGCAATTTGCAATTTGTATGCTGCATGGTTCTGTACG |
| SNP_IGA_758767 | GAAGGTGACCAAGTTCATGCTGGTTGAACTGGCAGCAAACAATGAT | GAAGGTCGGAGTCAACGGATTGTTGAACTGGCAGCAAACAATGAG | TTCGGATGGCGTCACGTTGTAGAA | CTTTATCATGTTGAGCGAATGCCTTTTTAAAATGATCTCCAGTTTGCTCTCGTACTTCGGATGGCGTCACGTTGTAGAACACGGGCAAAATCAATCTGGA[A/C]TCATTGTTTGCTGCCAGTTCAACCATTTTCGCAAGTTCATCCAGACACCATGTGGAGGTCGCATAGTTTGTTGAAAGAATGATGACTGAAATTTTCGATT |
| SNP_IGA_774557 | GAAGGTGACCAAGTTCATGCTGCACTATCAGGATCTGAGAAGATA | GAAGGTCGGAGTCAACGGATTGCACTATCAGGATCTGAGAAGATG | CTCTGCTGATGGTTTCGAGACTACAA | AGCACAATAGTAAAACTATGGCCACAATCAAGCACTTTTACAATCGGGGACAACAAAACTGAAAACAGAGTTAGCCTGCACTATCAGGATCTGAGAAGAT[A/G]TTAACATTCCCACACTTTGTAGTCTCGAAACCATCAGCAGAGGTTTGTTTAAGACTCAAAGGAACCTTCAGCTTGCAGTTGATGTTCCGTGACGGCGTGT |
| SNP_IGA_776826 | GAAGGTGACCAAGTTCATGCTGCCTCTGTAGCTTCCACCAAAT | GAAGGTCGGAGTCAACGGATTGCCTCTGTAGCTTCCACCAAAC | TTTGCTGATGATATTGCAGGCGACATTT | TCTACACTAGAAATTCTGGTATTCTTGATCACTAGAGGACATGGGAGAGCTCTTCTTTTGTTTGCTGATGATATTGCAGGCGACATTTCCATAGGGAAGC[A/G]TTTGGTGGAAGCTACAGAGGCACGCGGCAGAGGGGAAAAGAGATTCGGAGAAAGATGAGCAGAAGCAGTAACAATAGCCATTTGATAGATATCAGAGTAG |
| SNP_IGA_781003 | GAAGGTGACCAAGTTCATGCTGCACTTGTGAACCATCAGCCAA | GAAGGTCGGAGTCAACGGATTGCACTTGTGAACCATCAGCCAG | AACTGAATGGACATTGTGGACATGTCTTA | GTAAATTTAAATTATATAAGCAAGGGAAGAACCTTGAATGTTTTATAAGGGGTTGCATTTGATAAAACCATTGAAGTGTGCACTTGTGAACCATCAGCCA[A/G]CAAAACCTAGAATAAGACATGTCCACAATGTCCATTCAGTTAGATTTAAGTTTCAAAACTGCCTCATTTAGGAAAGCAAGAAGAAACCAACCCCGTTGAC |

| | | | | |
|---|---|---|---|---|
| SNP_IGA_784777 | GAAGGTGACCAAGTTCATGCTGTTATATACATCAAGAGAGGCACATACAA | GAAGGTCGGAGTCAACGGATTATATACATCAAGAGAGGCACATACAG | GGTTCCATTTTTTCTTCTTAATCTCCACAT | GATGGCACTGGAAATTTCACAAATGTGACGGATGCCGTGTTAGCAGCACCAGATTATAGCATGAGAAGATATGTTATATACATCAAGAGAGGCACATACA[A/G]AGAGAATGTGGAGATTAAGAAGAAAAAATGGAACCTAATGATGATTGGAGATGGTATGGATGCTACTATAATCTCTGGTAACAGAAGCTTTGTGGATGGC |
| SNP_IGA_794167 | GAAGGTGACCAAGTTCATGCTGAATATGCAGTTGGACACTCTGACT | GAAGGTCGGAGTCAACGGATTAATATGCAGTTGGACACTCTGACC | GAACCTCTTTGATAGGATCAGGCACTT | AGTATTTTAAGAACGGTGTCCCCAGGCATCCTGTTAACTTGTAGATGGCGAACCTCTTTGATAGGATCAGGCACTTCACTTTTCATTATTTCATTGTTCA[A/G]GTCAGAGTGTCCAACTGCATATTCAGGTTCCAATTCTTTAACAATATTATGATAAAATTCATCAACCTCAGGGGAGTCTGGGTTGGAGGACGTAGGTTTC |
| SNP_IGA_797492 | GAAGGTGACCAAGTTCATGCTAGGCCTCGTAACCAATCATTACTC | GAAGGTCGGAGTCAACGGATTATAGGCCTCGTAACCAATCATTACTA | GAGTTTGAAGATCGTTGGAAAGAGATGAT | AATCTATTAAATTTGCCTTGGAGAACATTGTGTATGATTCATTGACCAATATTGAGTTTGAAGATCGTTGGAAAGAGATGATTGAGAAGTATGAGTTACA[G/T]AGTAATGATTGGTTACGAGGCCTATATGATGAAAGACGTCGTTGGGTGCCAAGCTTTGTGAAAGGAAGTTTTTGGGCGGGCATGTCTACCACACAACGAA |
| SNP_IGA_803699 | GAAGGTGACCAAGTTCATGCTCACCACCTCCACTCTGATCTTGT | GAAGGTCGGAGTCAACGGATTACCACCTCCACTCTGATCTTGC | GGAAGGAAGGGAATCACAGATTCGAA | GTGAGTCTCATTATACAGTCCTTTTTTCCCTAGATACCACTGTTCAAGATGAGAATGAACTGGAAGGAAGGGAATCACAGATTCGAAAAGCTTTTGATGC[A/G]CAAGATCAGAGTGGAGGTGGTGGTTTTATTAGTGTGGAAGGCTTTCATCAAGTCCTTAAAGAAACTAATGTAGAACTTCCAACTGAGAAGGTTGACCTCC |
| SNP_Pp08-3702593 | GAAGGTGACCAAGTTCATGCTGGGATGTGCCGAGGCACG | GAAGGTCGGAGTCAACGGATTGTGGGATGTGCCGAGGCACA | GCAAGCTTTGTCTCGCCTTAAGCAT | CAACATCCCTTTTGCCTGCAGCACCCCCGTGCGTAGCCACATTGCCTTGTAACATGCAAGCTTTGTCTCGCCTTAAGCATCCCTTCACCTTCGGCACCCC[C/T]GTGCCTCGGCACATCCCACTGCGACATGCAAGCTTTGTGCATTTGACACCCCCGTTGCCTTCAGCACCCCCATGCCTAGCCACATTGCCTTACAACATGC |
| SNP_IGA_821894 | GAAGGTGACCAAGTTCATGCTTTTGAATGATTTCATTCACACCTCCC | GAAGGTCGGAGTCAACGGATTCATTTTGAATGATTTCATTCACACCTCCT | CATTATCATCTGGTTCGTTGGCAGGAT | TATACAAGTGTCCAGGCTCTCCGTTGATTGTGAAGGCATCTGAAACGTTTGGATCGCCACCTGTTGAGAGAGCATTTTGAATGATTTCATTCACACCTCC[C/T]TTGTACCAGGATCCTGCCAACGAACCAGATGATAATGAAAAATAATTATATGGGATTCGATGTTGCAACTGTATGATGTAACATTTTTTGTTTTCATGTC |
| SNP_IGA_851849 | GAAGGTGACCAAGTTCATGCTTCAATTGTTTGGGAGAACTTTATTAAG | GAAGGTCGGAGTCAACGGATTCCTTCAATTGTTTGGGAGAACTTTATTAAA | GCTACCATATGCATACACTTTATTGCAATA | GACTCAACAAGTTTGATGTCCCATTTTTTTTGCTACCATATGCATACACTTTATTGCAATATTTGCATTTTGCCTTAGGATTTTTTGGATCATCTTGCAT[C/T]TTAATAAAGTTCTCCCAAACAATTGAAGGAGGTCTAGCAGGTTTTCTCTAACCATCTTCAATGGGAGCAGAATCATTGGCTTTGGTTTCTTGTGGCAACA |

| | | | | |
|---|---|---|---|---|
| SNP_IGA_860815 | GAAGGTGACCAAGTTCATGCTTATTGTGCTACTTTTAGGTCAAACTTTCA | GAAGGTCGGAGTCAACGGATTGTGCTACTTTTAGGTCAAACTTTCG | GTTCTGCTGTCTTGCCATCAGCTTT | ATTACTGTCCAGCTCTTGTCAGAGATTGATACCTTTTCTATTACTAATCTGTCATTTTGTCTGATGTATTTTTATTGTGCTACTTTTAGGTCAAACTTTC[A/G]GAGAAAGCATTGTTGCTGGAAAAAGCTGATGGCAAGACAGCAGAACTTACAGCAAAGGTTAATGAACAACAAAAGTTAATCCAAAAGTTGGAGGATGACA |
| SNP_Pp08-13670362 | GAAGGTGACCAAGTTCATGCTGGGCTGCTTACAGGTGTTTTGTG | GAAGGTCGGAGTCAACGGATTAGGGCTGCTTACAGGTGTTTTGTA | ATGCTAAGCCAAAATCTTATCTGAAAGCAA | TAAGAAGCCTTATTTTCCCCTTTTGATGCTCAACTCATGCTGACCCTGCTCCCATACCACAAGCTCTTCATGCTAAGCCAAAATCTTATCTGAAAGCAAG[C/T]ACAAAACACCTGTAAGCAGCCCTCATTTTCATTGGTGAAGGTAACCAGAACACTTCCTAAGGTTTTGCTCCCCTTTCGAAGTGCTTTTGGGGCTTGATTT |
| SNP_IGA_871082 | GAAGGTGACCAAGTTCATGCTGTACAAGAAAGCAAACCCCAGCAA | GAAGGTCGGAGTCAACGGATTACAAGAAAGCAAACCCCAGCAG | CCAATCTTTGGAGATGGTTTACCAGAATA | TTTTCAGAGCTCTTCAAACAGTTCCCATTCTGATCAGTCTTCACTAAAGCTCTGGCTTTCTTCTTCCCTCTTAGTCTGTACAAGAAAGCAAACCCCAGCA[A/G]TGAAGCTATGACTGCACCAAAAATATTCTGGTAAACCATCTCCAAAGATTGGTGCAAAAAGGGTTAAAGAAGATTATGGTGGCACTCTCTGAATTTAAGT |
| SNP_IGA_878044 | GAAGGTGACCAAGTTCATGCTCAGTTGCAAGGGAGAACAGGGTA | GAAGGTCGGAGTCAACGGATTAGTTGCAAGGGAGAACAGGGTG | GGTTGTATACTCTGCTCACCGGAA | ACATCAAAGTTTGTTACTTTTGGATCAAATTATTTCTTGAAGTTACTGTAGGTAAAATTGGATGGGAATTTACCTGGACAGTTGCAAGGGAGAACAGGGT[A/G]ATTCCGATTTTCCGGTGAGCAGAGTATACAACCCCTTCGGACTCACTTCCGAGCTTAATTCCGGTAGCCCAGCCGGCGACCCCAATGGCATAACCAGATA |
| SNP_IGA_883524 | GAAGGTGACCAAGTTCATGCTAGAGGTCATTATACTGTGAAGGTGC | GAAGGTCGGAGTCAACGGATTGAGAGGTCATTATACTGTGAAGGTGA | CGCTTTCTCAAGGTTTTCCCATCTCTT | AACATTCTGTTTTCTCATTAATGTAGCCGCCCGATACATGTAAGGGTAAGTGAGAGTTGGGTCCAGCTCAGACGCTTTCTCAAGGTTTTCCCATCTCTTA[G/T]CACCTTCACAGTATAATGACCTCTCCTGATACATCCATCCAAGTGGGGTAAGAGAGCAGATTACAGAGCTCATCTTTTCATACGACCAAAGTTTATGGCC |
| SNP_IGA_884755 | GAAGGTGACCAAGTTCATGCTGTAGAAGGCAAGGAAGATATGGT | GAAGGTCGGAGTCAACGGATTGTAGAAGGCAAGGAAGATATGGC | TCACCTGCAACAGGTTGCTCATCAA | AGCCACCATCCCTTCAGCAGATGGAAGGCATCATGCTTCCACCAGTAGTCCATCTGCTTCACCTGCAACAGGTTGCTCATCAACAGTACCTGAAAAAGGT[A/G]CCATATCTTCCTTGCCTTCTACAGATACCAAGTTTTCCCAGAAATCTGGGTTGGAAGAGATATCTGGACCTAAGACCCCAGTTTGCACCTCCGAATCCCC |

**Table S3**. Heterozygous SNPs for ‘Sweet Dream’ and SNP density per chromosome

| Chromosome | #Heterozygous SNPs | Total chrom size (bp) | SNPs/Mb |
|---|---|---|---|
| Pp01 | 6,913 | 47,851,208 | 144.5 |
| Pp02 | 22,078 | 30,405,870 | 726.1 |
| Pp03 | 12,325 | 27,368,013 | 450.3 |
| Pp04 | 14,278 | 25,843,236 | 552.5 |
| Pp05 | 9,937 | 18,496,696 | 539.7 |
| Pp06 | 10,554 | 30,767,194 | 343.0 |
| Pp07 | 11,701 | 22,388,614 | 522.7 |
| Pp08 | 6,665 | 22,573,980 | 295.3 |
| Total | 94,451 | 225,694,811 | 418.7 |

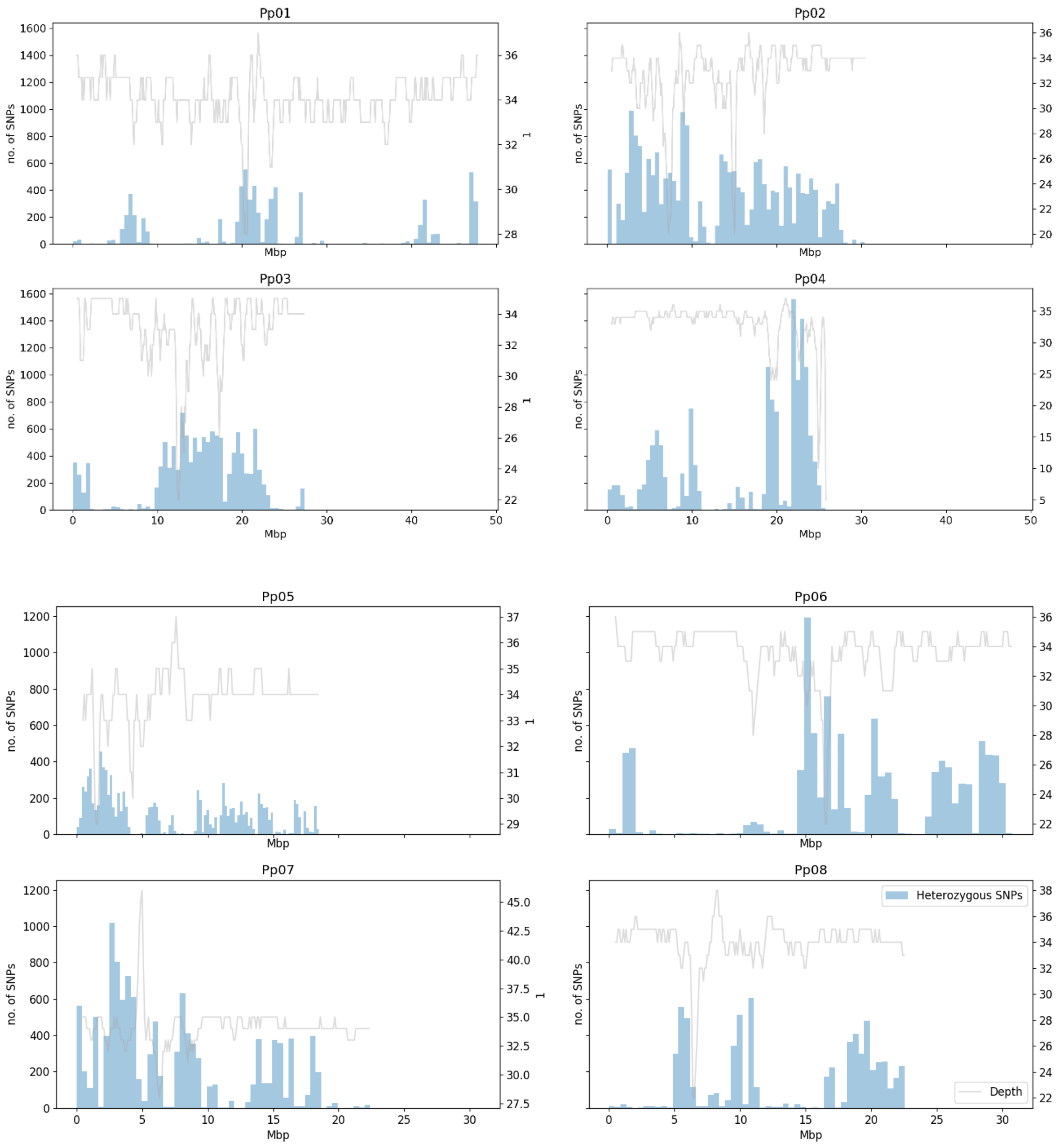


**Figure S1**. SNP and depth distribution in ’Sweet Dream’. The distribution of heterozygous SNPs is shown across the eight chromosomes of the peach genome in bins of 500kbp. Grey line represents median read coverage across chromosome, calculated in windows of 500Kb with a step of 100kb.

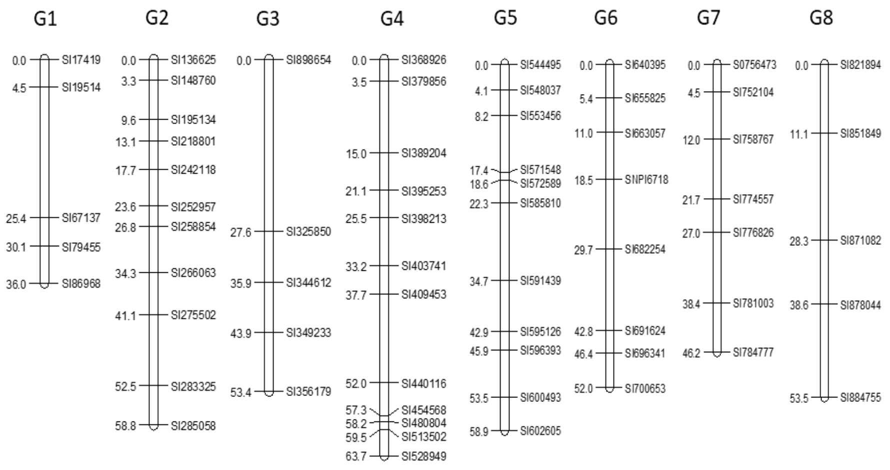


**Figure S2.** Linkage map of peach ‘Sweet Dream’ with 64 SNPs.
